# Synaptic activity controls local exposure of an ‘eat-me’ signal via ANO3-ITPR1 signaling

**DOI:** 10.64898/2026.06.26.734915

**Authors:** Kunal Shroff, Molly C. O’Brien, Tyler J. Smith, Jialing Fang, Kelly S. Yu, Sonia I. Lombroso, Shengjie Feng, Chao Chen, Gregory A. Mohl, Indigo V.L. Rose, Nidhi R. Lokesh, Mark E. Pownall, Lily Y. Jan, Martin Kampmann

## Abstract

Neuronal synapses are eliminated during brain development and disease through pruning by glial cells. Individual synapses are marked for engulfment by ‘eat-me’ signals, which include externalized phosphatidylserine. In apoptotic cells, phosphatidylserine externalization is driven by caspase-dependent activation of Xkr scramblases^1^ and inactivation of specific flippases^2^. Localized caspase activation at neuronal synapses can mediate spatially-restricted synaptic phosphatidylserine exposure leading to synapse pruning by glial cells during development and neurodegeneration^3–6^. It is unknown which caspase-regulated flippases and scramblases promote synaptic phosphatidylserine exposure and whether there are any caspase-independent mechanisms of phosphatidylserine exposure relevant for synapse elimination. To address this question, we here develop a scalable CRISPR screening approach, COMPASS-seq (compartment-anchored sgRNA screen sequencing), to uncover the genetic underpinnings of subcellular phenotypes. COMPASS-seq is compatible with *in vitro* and *in vivo* systems and a wide range of subcellular compartments; we here apply it to the neuronal synapse. We discover that inhibition of the caspase-independent, calcium-activated anoctamin ANO3 (TMEM16C) is sufficient to increase synapse numbers *in vivo*. ANO3 co-localizes with IP3 receptor 1 (ITPR1), a calcium channel in the endoplasmic reticulum, to form a postsynaptic signaling platform that drives spatially restricted phosphatidylserine exposure at synapses. Activation of the ITPR1 calcium channel activity is sufficient to drive synaptic phosphatidylserine exposure via an ANO3-dependent but caspase-independent mechanism. Our results suggest a mechanism for integrating synaptic activity information to control synaptic pruning. The role of ANO3 in regulating synapses could shed light on the mechanisms underlying its numerous associations with both dementia^7^ and other neurological diseases^8,9^.

## Main text

Synaptic activity plays a critical role in the proper maturation of neuronal circuits across the brain during postnatal development^10–13^. This generally involves an initial period of rapid synapse formation followed by a prolonged window of brain-wide, experience-driven synapse loss^10,11,13,14^. Some portion of this experience-dependent synapse loss is mediated by the neuron-intrinsic mechanism of long-term depression, which is classically driven by both NMDA receptor signaling and signaling from the Type I metabotropic glutamate receptor (mGluR1) to the IP3 receptor (ITPR1) ^15–17^. These receptors directly link synaptic activity patterns to synaptic calcium changes that are thought to drive synapse weakening and loss through various neuron-intrinsic mechanisms.

Beyond these well-characterized neuron-intrinsic mechanisms of synapse loss, recent work has uncovered a role for glial cells in mediating synapse loss during development^18–20^. Glial cells have been proposed to selectively engulf synapses tagged with components of the complement system^18,19^. Several studies have shown that complement binds to exposed phosphatidylserine (PtdSer) at synapses^21,22^. PtdSer itself can also directly bind to several glial receptors that mediate synapse loss during development^23–25^. Classically, PtdSer serves as a signal for apoptotic cell engulfment by macrophages and is regulated by apoptotic caspase activity^26^. Recent results suggest that caspase activity can regulate glia-mediated synapse loss during development^5,6^. However, PtdSer exposure can also be regulated by several families of caspase-independent flippases and scramblases^27^. It is unknown which specific caspase-dependent flippases and scramblases promote PtdSer exposure at synapses and whether there are any caspase-independent mechanisms of PtdSer exposure engaged at neuronal synapses.

To uncover regulators of synaptic numbers, we here develop a technology to enable CRISPR screens based on phenotypes in subcellular compartment, COMPASS-seq (<u>com</u>partment-<u>a</u>nchored <u>s</u>gRNA <u>s</u>creen <u>seq</u>uencing), and integrate it with a synapse-targeted trafficking protein. Using this technology, we discover a role for ANO3 (TMEM16C), a calcium-dependent scramblase^28^, which drives PtdSer exposure upon activation. Within neurons, we find that ANO3 selectively drives PtdSer exposure at synapses. ANO3 colocalizes postsynaptically with ITPR1, and ITPR1 activity can activate ANO3 to drive synaptic PtdSer exposure. Our findings delineate a caspase-independent mechanism by which information on local synaptic activity is integrated by a spatially restricted signaling platform to selectively tag synapses for elimination by glial cells.

## Results

### *In vivo* CRISPR screening platform for synaptic phenotypes

Pooled CRISPR-based screens have emerged as powerful tools to uncover the individual contribution of thousands of genes to a phenotype of interest within a single experiment^29,30^. They are enabled by introducing pooled libraries of single guide RNAs (sgRNAs) into populations of cells to perturb different genes in individual cells^29,30^. One of the principal issues when designing any novel pooled CRISPR screen is how to identify the genetic perturbations causing a phenotype of interest. Classically, this linkage of phenotypes to genetic perturbations is achieved by amplifying and sequencing the sgRNA-expressing element integrated into the genome of cells displaying the phenotype of interest^30^. However, this strategy restricts the scope of phenotypes that can be investigated with pooled CRISPR screens to those readily associated with the nucleus harboring the genome. This restriction creates a substantial barrier to the application of pooled CRISPR screens to the study of phenotypes in distal subcellular compartments, such as the synapse. Synapses are distant from the nucleus, making it difficult to maintain physical linkage between the nucleus, which harbors the sgRNA, and the synapse manifesting the phenotype.

To enable pooled CRISPR screens for subcellular phenotypes, with a first application to synaptic phenotypes in neurons, we devised a system to localize a copy of the sgRNA to the synapse, enabling the linkage of synaptic phenotypes to information on the gene perturbed in the nucleus (Figure 1a). The sgRNA sequence and associated U6 promoter is placed within the 3’ untranslated region of an mRNA transcript driven by the neuron-specific hSYN promoter. We will refer to this mRNA transcript containing an sgRNA sequence within its 3’ untranslated region as the guidecode.

**Figure 1.**
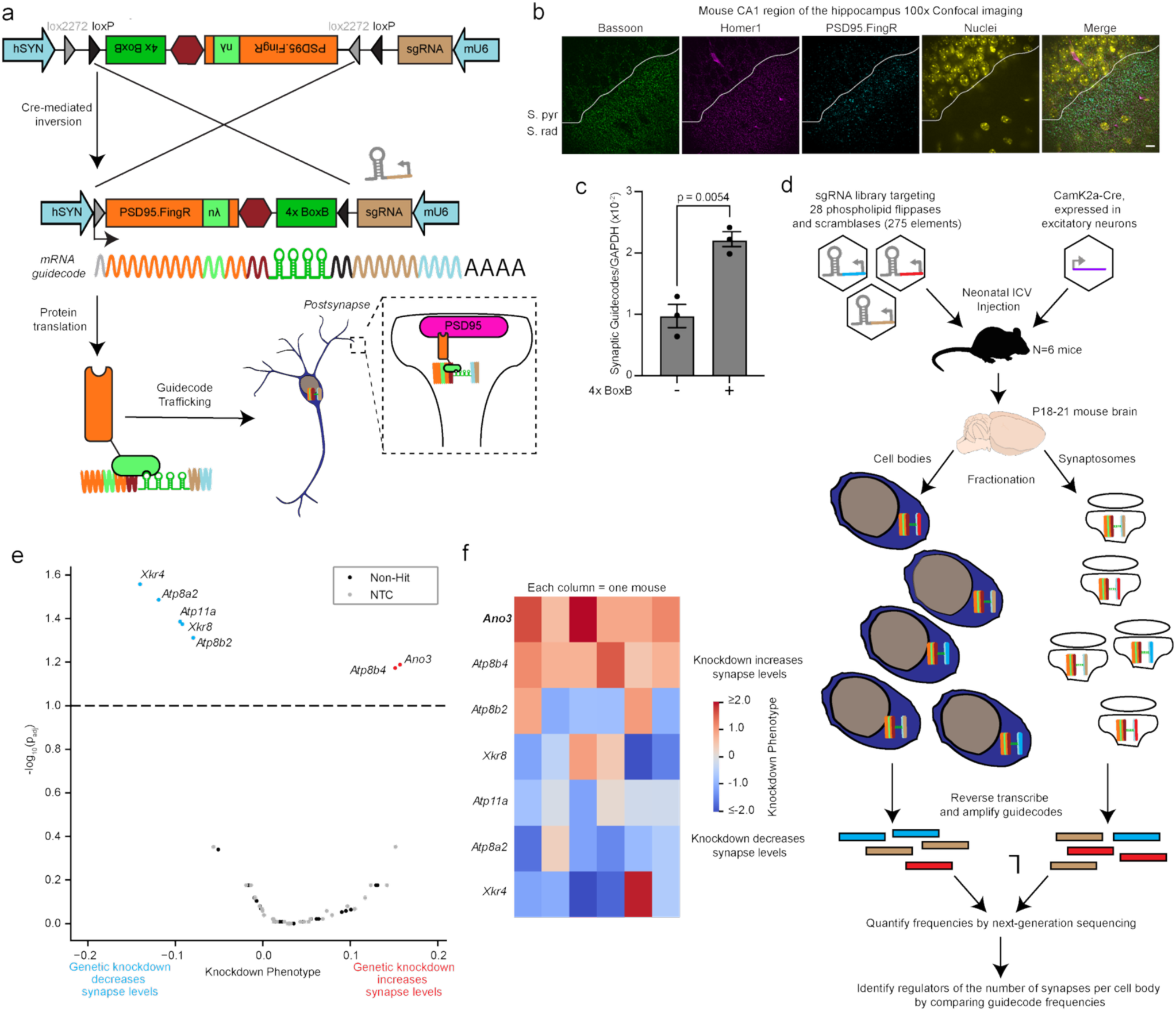
Synaptic screening platform uncovers Ano3 as a postsynaptic modifier of synapse numbers in mouse brains. (a) AAV plasmid for cell-type-specific *in vivo* CRISPR screens for factors controlling synapse levels. An mU6-sgRNA element is incorporated within the 3’ UTR of an hSYN-dependent transcript hereafter referred to as a guidecode. The construct includes a Cre-On flip excision system such that when Cre recombinase is expressed, a portion is irreversibly inverted, enabling (1) cell-type specific protein expression of a synaptic trafficker consisting of a modified PSD95.FingR construct fused to an RNA-binding nλ domain and (2) cell type-specific incorporation of the BoxB structured RNA into the guidecode. The BoxB RNA structure selectively binds to the nλ domain, enabling targeting of the guidecode to the postsynaptic density. (b) Immunofluorescent staining for the synaptic trafficker alongside presynaptic (Bassoon) and postsynaptic (Homer1) markers in CA1 hippocampus of P28 mice co-injected with AAV-PHP.eB-packaged pTS026 and CamK2a-Cre at birth (Scale bar = 10 μm). (c) Digital PCR to quantify the efficacy of the nλ-BoxB system to traffic mRNA into the synaptosome fraction by comparing synaptosome guidecode levels in mice injected with trafficking constructs in which the 4xBoxB domain is either present or deleted from the guidecode (mean ± s.e.m., n = 3 mice per construct, p-value calculated by two-sided unpaired t-test). (d) Workflow for an *in vivo* CRISPR screen for postsynaptic factors in excitatory neurons affecting synapse numbers. A 275-element sgRNA library targeting 28 phospholipid flippase and scramblase genes was cloned into pTS026 and packaged into AAV (PHP.EB capsid) alongside CamK2a-Cre. These AAVs were co-injected into 6 neonatal LSL-CRISPRi mice. Brains were harvested on P18-21 and fractionated into somatic and synaptosome fractions. Guidecode within these fractions were extracted and selectively amplified by RT-PCR. (e) Results from the screen. Hit genes (FDR threshold = 0.1) are annotated. (f) Knockdown phenotype for each hit gene from each individual mouse (columns).

To help ensure that the guidecode is well represented at synapses distal to the nucleus, we integrated genetic elements for trafficking mRNA barcodes along neuronal projections with new nanobody tools for labeling synapses (Figure 1a). Specifically, we fused the PSD95.FingR construct, which binds to the endogenous postsynaptic protein PSD-95^31^, to the nλ peptide, which binds an RNA stem loop motif, BoxB^32^. This RNA motif was incorporated into the 3’ untranslated region of the guidecode. This strategy anchors guidecodes to endogenous PSD95 to enable the identification of CRISPR-based perturbations at the synapse and thereby provide a basis for pooled CRISPR screens for synaptic phenotypes. We refer to this <u>comp</u>artment-<u>a</u>nchored <u>s</u>gRNA <u>s</u>creen <u>seq</u>uencing approach as COMPASS-seq.

To facilitate neuronal subtype-specific *in vivo* screens based on our previous CrAAVe-seq strategy^33^, we generated an AAV vector, pTS026, which contains COMPASS-seq technology within a Cre-dependent flip-excision genetic element (Figure 1a). To test the system, AAVs harboring pTS026 and CamK2a promoter-driven Cre recombinase (CamK2a-Cre) were introduced into neonatal mice by intracerebroventricular (ICV) injection. The genotype, age, sex, and injection information for each mouse used in this study is provided in Supplementary Table 1. The nanobody formed distinct puncta within the synapse-rich *stratum radiatum* layer of the hippocampus, colocalizing with synaptic markers Bassoon and Homer1 (Figure 1b). This finding suggests that our nλ peptide fusion does not interfere with the selectivity of PSD95.FingR nanobody labeling. We prepared synaptosomes from the brains, extracted total RNA, converted to cDNA, and performed a digital PCR to quantify the number of guidecodes within the synaptic fraction of the brain. Guidecodes derived from the pTS026 vector were robustly detected and enriched relative to a control construct lacking the 4x BoxB element (Figure 1c). RNA sequencing of the synaptosome and cell body fractions confirmed that our synaptosome fraction is broadly enriched for synaptic mRNAs (Extended Data Figure 1, Supplementary Tables 4-6).

### *In vivo* CRISPR screen for synapse numbers

To uncover mechanisms affecting synapse numbers through the exposure of the eat-me signal PtdSer^22^, we cloned a library of 275 sgRNAs targeting all 28 known mouse phospholipid flippases and scramblases into pTS026 (Figure 1d). Protospacer sequences are provided in Supplementary Table 7. We introduced AAVs harboring this library and CamK2a-Cre into six neonatal mice by ICV injection. To maximize the effects of developmental glia-mediated synapse elimination, we collected the brains 18-21 days after birth, which is a period characterized by enhanced glia-mediated synapse loss across many brain regions^11,17^. Each brain was fractionated into cell bodies and synaptosomes. We extracted the guidecodes from each fraction, amplified the sgRNA sequences from cells expressing Cre using reverse-transcriptase PCR, and quantified the differential frequency of sgRNA sequences between the cell body and synaptosome fractions using next-generation sequencing to identify genetic regulators of synapse numbers. Given that the trafficker used for this screen targeted postsynaptic PSD95, this screen is expected to predominantly identify postsynaptic cell-autonomous regulators of synapse numbers during development.

The screen uncovered 5 genes knockdown of which decreased synapse numbers and 2 genes knockdown of which increased synapse numbers (p_adj_ < 0.1) (Figure 1e, Supplementary Table 8). Effect sizes were consistent across individual mice (Figure 1f, Supplementary Table 9). Among the hits, we identified a single scramblase, Ano3, whose knockdown increased synapse numbers (Figure 1e). Given that knockdown of a scramblase would be predicted to decrease PtdSer exposure^1^, we hypothesized that Ano3 may be a key driver of PtdSer exposure specifically at synapses, which could explain its effect on synapse numbers.

### ANO3 regulates synaptic phosphatidylserine exposure

To investigate the molecular mechanism by which ANO3 affects synapse numbers, we optimized an *in vitro* human iPSC-derived neuron-astrocyte coculture system to recapitulate robust synapse formation and activity (Extended Data Figure 2). Neuronal knockdown of ANO3 in this system appeared to substantially decrease exposed PtdSer at synapses (Figure 2a, Extended Data Figure 3a). To quantify this phenotype more rigorously, we developed an FM dye-based synaptosome flow cytometry assay to rapidly quantify PtdSer exposure across thousands of active synaptosomes per sample (Extended Data Figures 4, 5). We found that changes in detected synaptic PtdSer exposure were largely independent of changes in synaptosome volume (Extended Data Figure 5k). We validated our synaptosome flow cytometry assay and our biological system by confirming the effects of two previously reported regulators of synaptic PtdSer exposure: amyloid-β oligomers, which has been shown to increase synaptic PtdSer exposure^25,34^, and pan-caspase inhibitor Z-VAD-FMK, which has been shown to decrease synaptic PtdSer exposure^6^. Amyloid-β oligomer treatment increased synaptic PtdSer exposure by ∼100% (Extended Data Figure 4e-f), while increasing synaptosome volume by ∼7% (Extended Data Figure 5a). Meanwhile, Z-VAD-FMK treatment decreased synaptic PtdSer exposure by ∼50% (Extended Data Figure 4g-h), and had no effect on synaptosome volume (Extended Data Figure 5b). Neither of these treatments affected somatic PtdSer exposure (Extended Data Figure f,h).

**Figure 2.**
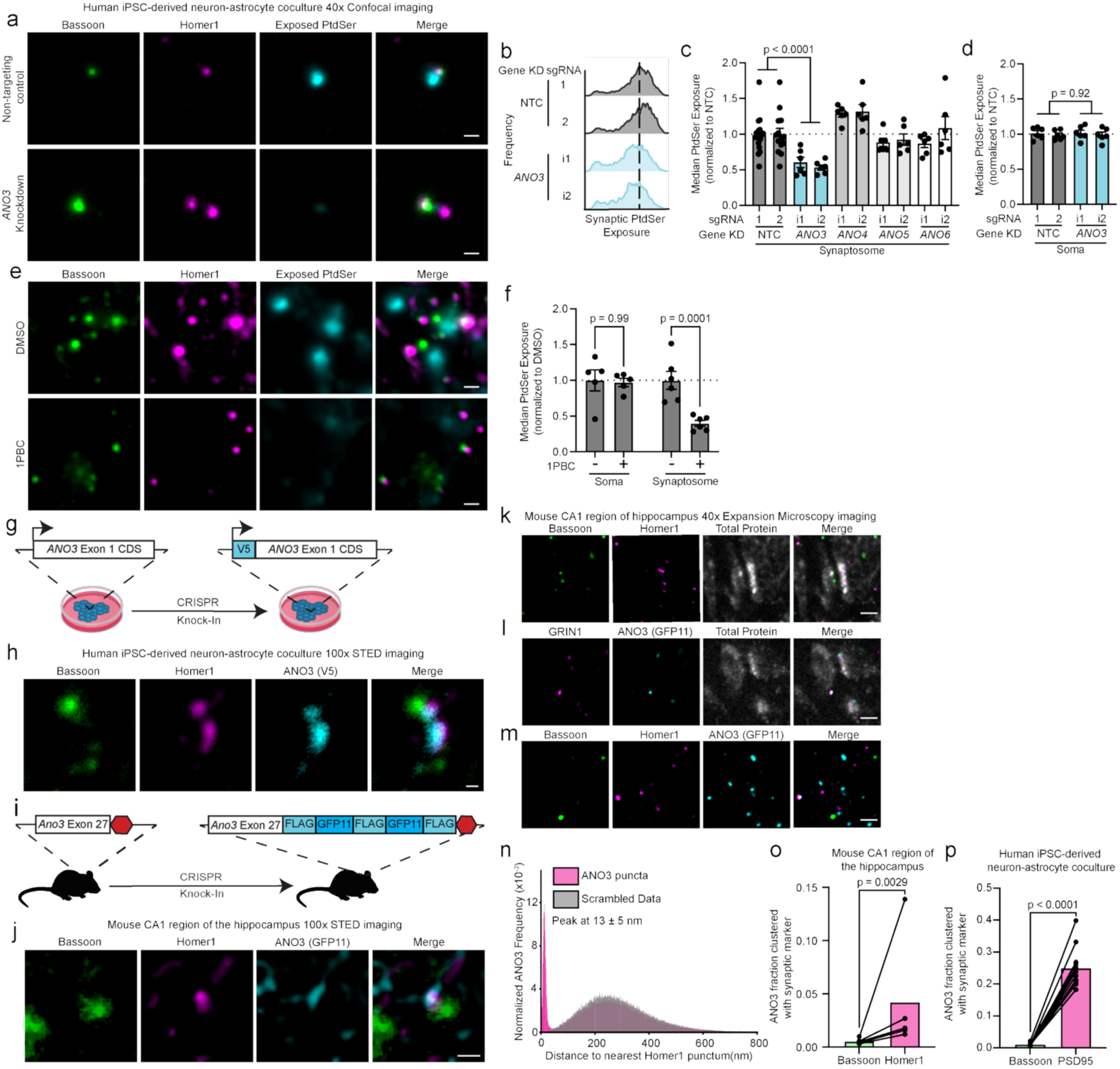
ANO3 is postsynaptically localized and its inhibition reduces externalized phosphatidylserine at the synapse, but not the soma. (a) CRISPRi-mediated knockdown of ANO3 within human iPSC-derived neurons drives loss of synaptic PtdSer exposure relative to a non-targeting control sgRNA, as visualized by immunofluorescent staining for presynaptic Bassoon, postsynaptic Homer1, and exposed phosphatidylserine (PtdSer) via pre-fixation Annexin-V staining in human iPSC-derived neurons cocultured with human iPSC-derived astrocytes for 21 days. Representative images are shown here (Scale bar = 1 μm); large fields of view are available as source data. (b-c) Quantification of synaptic PtdSer exposure via synaptosome flow cytometry using synaptosomes derived from neurons where either one of the neuronally expressed anoctamins with scramblase activity is knocked down or a non-targeting control sgRNA (NTC) is expressed. (b) Representative histograms of synaptic PtdSer exposure generated by synaptosome flow cytometry (c) Quantification of synaptic PtdSer exposure generated by synaptosome flow cytometry (mean ± s.e.m., n=6 wells, p-values calculated by one-way unpaired ANOVA followed by two-sided unpaired t-tests comparing against the NTCs with a Holm-Šídák correction for multiple comparisons applied) (d) Flow cytometry of neuronal cell bodies reveals that ANO3 knockdown has no effect on somatic PtdSer exposure (mean ± s.e.m., n=6 wells, p-values calculated by two-sided unpaired t-test). (e-f) 72-hour treatment with 50 μM 1PBC, a pharmacological inhibitor of anoctamins, including ANO3, similarly reduces synaptic PtdSer exposure as visualized by immunofluorescent staining for presynaptic (Bassoon), postsynaptic (Homer1), and exposed phosphatidylserine (PS) in human iPSC-derived neuron-astrocyte cocultures. (f) Quantification of synaptic and somatic PtdSer exposure upon 72-hour treatment with 50 μM 1PBC treatment in human iPSC-derived neuron-astrocyte cocultures shows that treatment selectively reduces synaptic PtdSer exposure with no effect on somatic PtdSer exposure (mean ± s.e.m., n=5-6 wells, p-values calculated by one-way unpaired ANOVA followed by two-sided unpaired t-tests comparing against DMSO control with a Holm-Šídák correction for multiple comparisons applied). (g) Schematic of the V5-tag incorporated onto the N-terminus of endogenous ANO3 within human iPSCs. (h) Immunofluorescent staining and imaging with super-resolution STED microscopy reveals that ANO3(V5) colocalizes to a greater extent with the postsynaptic marker Homer1 than with the presynaptic marker Bassoon in human iPSC-derived neuron-astrocyte cocultures. Representative images are shown here (Scale bar = 200 nm); large fields of view are in Extended Data Figure 7b. (i) Schematic of the FLAG-GFP11 tag incorporated at the C-terminus of endogenous ANO3 in mice. (j) Immunofluorescent staining and imaging with super-resolution STED microscopy reveals that ANO3(GFP11) colocalizes to a greater extent with the postsynaptic marker Homer1than with the presynaptic marker Bassoon in CA1 hippocampus of adult mice. Representative images are shown here; large fields of view are in Extended Data Figure 7a (k-l) Expansion microscopy pan-staining confirms that ANO3 localizes to the postsynaptic density of CA1 hippocampal synapses in *stratum radiatum.* (k) Representative staining of presynaptic marker Bassoon, postsynaptic marker Homer1, and a total protein stain to visualize the synaptic ultrastructure (Scale bar = 200 nm). (l) Representative staining of postsynaptic marker GRIN1, GFP11-tagged ANO3, and a total protein stain to visualize the synaptic ultrastructure (Scale bar = 200 nm). (m) Representative triple stain of presynaptic marker Bassoon, postsynaptic marker Homer1, and GFP11-tagged ANO3 on ∼20x expanded CA1 region of the hippocampus from P28 mice (Scale bar = 200 nm). (n) Histogram of nearest-neighbor analysis of the triple stain expanded tissue showcases the fraction of ANO3 puncta that cluster with Homer1 beyond random chance. (o) Quantification of the fraction of ANO3 puncta that cluster with either Bassoon or Homer1 in mouse CA1 hippocampus (n=5 fields of view across 3 mice collected across 2 sample processing runs, p-value calculated by a two-sided paired ratio t-test). (p) Quantification of the fraction of ANO3 puncta that cluster with either Bassoon or PSD95 in human iPSC-derived neuron-astrocyte cocultures (mean ± s.e.m., n=12 fields of view across 3 wells collected across 3 sample processing runs, p-value calculated by two-sided paired ratio t-test).

Knockdown of ANO3 reduced synaptic PtdSer exposure by ∼50%, whereas targeting all other neuronally-expressed, plasma membrane-associated, anoctamin scramblases (ANO4, ANO5, ANO6)^28^ did not decrease synaptic PtdSer exposure (Figure 2b-c, Extended Data Figure 3b-d). ANO3 knockdown did not affect synaptosome volume (Extended Data Figure 5c). In addition, ANO3 knockdown had no effect on somatic PtdSer exposure, aligning with our screen data, which indicated that ANO3 may selectively act at synapses rather than across the entire cell (Figure 2d). To exclude the possibility that the phenotype of ANO3 knockdown is driven by effects on neuronal maturation, we tested the effect of short-time pharmacological anoctamin inhibition. Treatment of cultures on day 18 with 1PBC, a pan-anoctamin inhibitor^35^, for three days reduced synaptic PtdSer exposure by ∼60% with no effect on synaptosome volume or somatic PtdSer exposure, recapitulating the effect of ANO3 knockdown (Figure 2e-f, Extended Data Figure 5d).

### ANO3 localizes to the postsynaptic membrane

Given that ANO3 emerged from the *in vivo* screen as a postsynaptic regulator of synapse numbers (Figure 1e), we hypothesized that ANO3 may be localized to the postsynaptic membrane and locally influence synaptic PtdSer exposure. To test this hypothesis, we used a CRISPR knock-in approach to tag the N-terminus of endogenous ANO3 with a V5 tag in iPSCs (Figure 2g, Extended Data Figure 6a). Confocal imaging revealed widespread ANO3 localization to synapses, although substantial amounts of ANO3 also localized to the cell body and neuronal projections (Extended Data Figure 7b). Using super-resolution STED microscopy, we observed that synaptic ANO3 selectively colocalized with the postsynaptic marker Homer1 relative to the presynaptic marker Bassoon within our human iPSC-derived neuron-astrocyte coculture system (Figure 2h). We further validated this finding by verifying the same result when using an endogenous antibody against endogenous ANO3 rather than the V5 tag (Extended Data Figure 8a,e).

To assess the localization of ANO3 *in vivo*, we endogenously tagged the C-terminus of ANO3 within mice with a FLAG-GFP11-FLAG-GFP11-FLAG tag (Figure 2i, Extended Data Figure 6b).

Mirroring results in iPSC-derived neurons, confocal microscopy revealed ANO3 localized to synapses as well as neuronal cell bodies and projections (Extended Data Figures 7a, 8d). STED microscopy revealed enhanced colocalization of ANO3 with the postsynaptic marker Homer1 relative to the presynaptic marker Bassoon within the CA1 region of adult mouse hippocampus (Figure 2j).

To assess the spatial localization of ANO3 relative to the pre- and postsynaptic membrane more quantitatively, we turned to expansion microscopy^36^. Specifically, we used pan-ExM-t^37^ to quantitatively assess the localization of ANO3 relative to pre- and postsynaptic markers across thousands of synapses. Our samples expanded by a linear factor of 21.6 ± 3.0-fold (mean ± standard deviation, n=250 synaptic profiles across 10 fields of view from 3 animals collected over 4 separate experiments), which corresponds to a <15-nanometer resolution when imaging on a standard confocal microscope (Extended Data Figure 9a-b, Supplementary Table 11).

To further verify the integrity of our expansion process and synaptic stains, we validated that presynaptic marker Bassoon and postsynaptic marker Homer1 localize on opposite sides of the synaptic cleft when staining expanded samples (Figure 2k). Furthermore, we found that staining for ANO3 and postsynaptic marker GRIN1 alongside the synaptic ultrastructure highlighted the localization of ANO3 to the outer edge of the postsynaptic density in the CA1 region of adult mouse hippocampus (Figure 2l).

To quantitatively assess the localization of ANO3 relative to pre- and postsynaptic markers, we stained expanded tissue samples with Bassoon, Homer1, and ANO3 and performed a nearest-neighbor analysis to quantify the degree of non-random clustering between ANO3 and each of the two synaptic markers (Figure 2m-n). This analysis quantitatively revealed that ANO3 preferentially clusters with the postsynaptic marker Homer1 relative to the presynaptic marker Bassoon *in vivo* (Figure 2o, Supplementary Table 12). These results were replicated when performed on our human iPSC-derived neuron-astrocyte cocultures using both the V5 tag as well as the endogenous ANO3 antibody to stain for ANO3 puncta (Figure 2p, Extended Data Figure 8b-c, Supplementary Table 12).

### ANO3 colocalizes with ITPR1 at the postsynaptic compartment

Given that ANO3 is a calcium-dependent scramblase^28^ localized to the postsynaptic membrane, we hypothesized that ANO3 may be activated by postsynaptic calcium sources. ANO1, a non-scramblase homolog^28^ of ANO3, has previously been reported to interact physically with the IP3 receptor ITPR1, an IP3-dependent endoplasmic reticulum calcium channel, within peripheral sensory neurons^38^. ITPR1 partially localizes to the postsynaptic compartment and has been functionally associated with mechanisms of synaptic long-term depression^39,40^. Therefore, we hypothesized that ANO3 may interact with ITPR1 to drive synaptic PtdSer exposure at neuronal synapses.

We observed that ITPR1 localizes throughout neurons, with substantial amounts in both the soma and at synapses across both of our experimental systems (Extended Data Figure 7c-d). STED microscopy further revealed that, at the synapse, ITPR1 preferentially localizes within the postsynaptic compartment in both human iPSC-derived neuron-astrocyte cocultures and the CA1 region of adult mouse hippocampus (Figure 3a,d). Finally, we used expansion microscopy alongside nearest neighbor analysis to quantitatively confirm that ITPR1 preferentially clusters with postsynaptic markers relative to presynaptic markers (Figure 3g-h, Supplementary Table 12).

**Figure 3.**
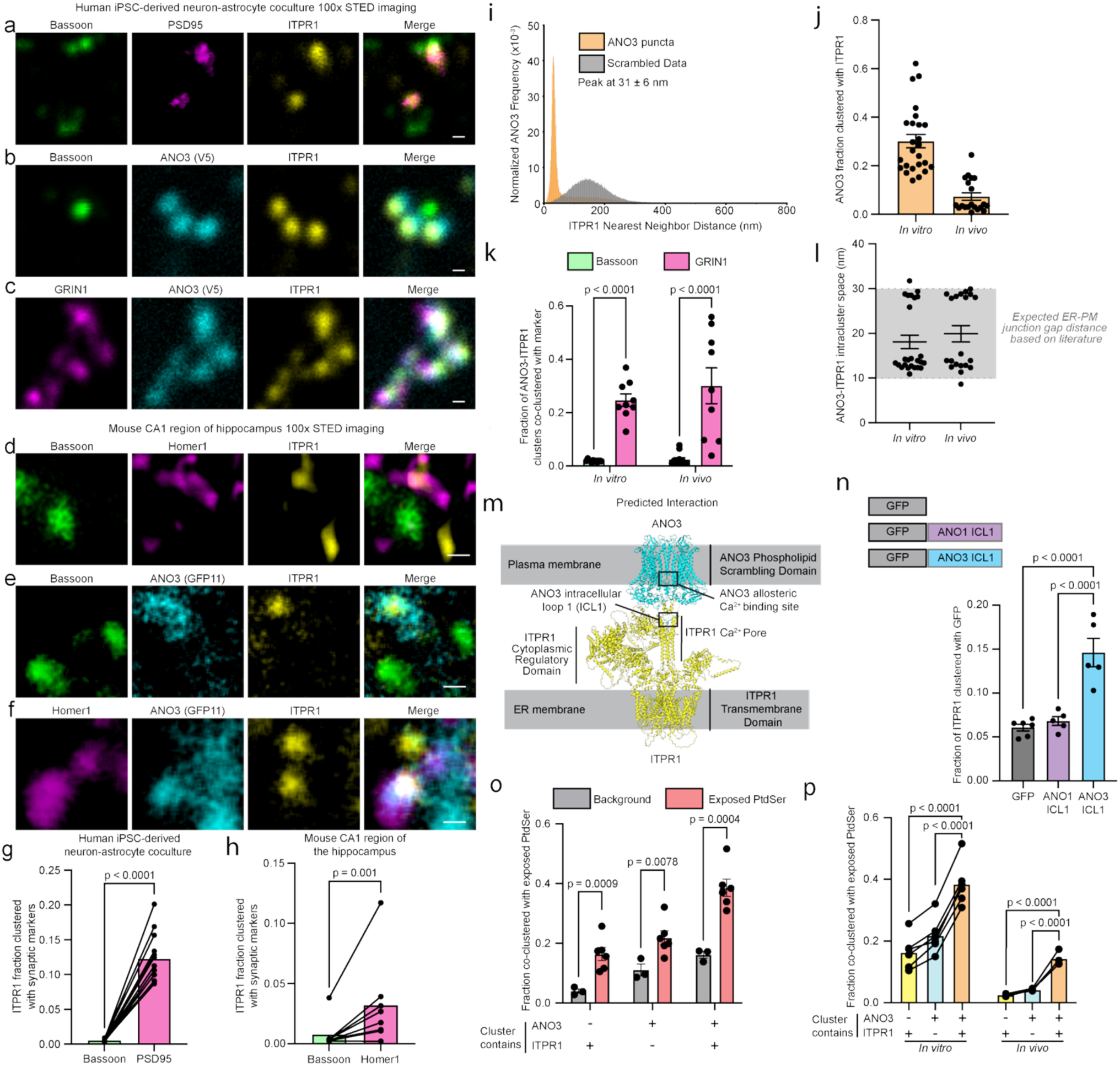
ANO3 co-localizes with the IP3 receptor in the postsynaptic compartment. (a-f) 100x STED imaging reveals that ANO3 colocalizes with ITPR1 (the neuronal IP3 receptor) at the postsynapse visualized by PSD95 and/or GRIN1 (a,b,c) in human iPSC-derived neuron-astrocyte cocultures and by Homer1 (d,e,f) in the adult mouse CA1 region of the hippocampus. Representative images are shown here (Scale bar = 200 nm); large fields of view are in Extended Data Figure 7c-h. (g-h) Nearest-neighbor analysis of ∼20x expanded samples confirms that ITPR1 predominantly clusters with postsynaptic markers (PSD95 *in vitro*, Homer1 *in vivo*) compared to presynaptic Bassoon in both (g) human iPSC-derived neuron-astrocyte cocultures (mean ± s.e.m., n=13 fields of view across 4 wells and 4 sample processing runs, p-value calculated by two-sided paired ratio t-test) and (h) adult mouse CA1 region of the hippocampus (n=7 fields of view across 3 mice and 3 sample processing runs, p-value calculated by two-sided paired ratio t-test). (i) Histogram of nearest-neighbor analysis of the ∼20x expanded samples showcases the fraction of ANO3 puncta that colocalize with ITPR1 beyond random chance. (j) Quantification of nearest-neighbor analysis of ∼20x expanded samples shows that ∼30% of ANO3 clusters with ITPR1 in human iPSC-derived neuron-astrocyte cocultures (mean ± s.e.m., n=15 fields of view across 6 wells and 4 sample processing runs) and ∼10% of ANO3 clusters with ITPR1 in mouse CA1 region of the hippocampus (mean ± s.e.m., n = 10 fields of view across 5 mice and 4 sample processing runs). (k) Nearest-neighbor analysis shows that ANO3-ITPR1 clusters are predominantly co-clustered with postsynaptic marker GRIN1 relative to presynaptic marker Bassoon (mean ± s.e.m., *In vitro*, n= 9 fields of view for GRIN1 localization and 12 fields of view for Bassoon localization across 3 wells and 3 sample processing runs. *In vivo*, n = 9 fields of view for GRIN1 localization and 11 fields of view for Bassoon localization across 3 mice and 2 sample processing runs, p-values calculated by two-sided unpaired lognormal t-test with a Holm-Šídák correction for multiple comparisons applied). (l) Expansion microscopy reveals that in both *in vitro* and *in vivo* samples, ANO3-ITPR1 clusters are on average separated by ∼10-30nm, which is consistent with the ER-PM junction gap distance in mammalian cells^41–43^. (m) AlphaFold modeling predicts that the first intracellular loop of ANO3 binds to the cytoplasmic C-terminal calcium channel domain of the IP3 receptor. This would place the ITPR1 calcium channel pore proximal to an allosteric calcium-binding site within the ANO3 homodimer. (n) (Top) Schematic of superfolder GFP control or superfolder GFP fused to either the first intracellular loop of ANO1 or the first intracellular loop of ANO3 fusion constructs. (Bottom) Expansion microscopy nearest-neighbor analysis reveals that the first intracellular loop of ANO3 enhances binding of superfolder GFP to ITPR1 within human iPSC-derived neuron-astrocyte cocultures (mean ± s.e.m., n= 5-6 fields of view from 1 well per condition across 2 sample processing runs, p-values calculated by one-way unpaired lognormal ANOVA followed by two-sided unpaired lognormal t-tests with a Holm-Šídák correction for multiple comparisons applied). (o) Nearest-neighbor analysis shows that both ANO3 puncta and ANO3-ITPR1 clusters co-cluster significantly with exposed PtdSer stained via pre-fixation Annexin-V staining in human iPSC-derived neuron-astrocyte cocultures (mean ± s.e.m., n = 3-6 fields of view across two wells from 1 sample processing run. p-values calculated by two-sided unpaired lognormal t-tests with a Holm-Šídák correction for multiple comparisons applied). (p) Nearest neighbor analysis shows that ANO3-ITPR1 clusters preferentially co-cluster with exposed PtdSer relative to independent ANO3 and independent ITPR1 clusters both *in vitro* and *in vivo* (mean ± s.e.m., *In vitro*, n = 6 fields of view across 2 wells and 1 sample processing run. *In vivo*, n = 4 fields of view across 1 mouse and 1 sample processing run, p-values calculated by one-way paired lognormal ANOVA followed by two-sided paired lognormal t-tests with a Holm-Šídák correction for multiple comparisons applied).

ITPR1 and ANO3 colocalized at postsynaptic compartments in both human iPSC-derived neuron-astrocyte cocultures as well as the CA1 region of adult mouse hippocampus (Figure 3b-c, 3e-f, Extended Data Figure 7e-h). ITPR1 puncta and ANO3 puncta clustered across both experimental systems (Figure 3i-j, Supplementary Table 13). These ANO3-ITPR1 clusters preferentially co-clustered with postsynaptic GRIN1 relative to presynaptic Bassoon, consistent with our earlier observations (Figure 3b,c,e,f,k; Extended Data Figure 7e-h, Supplementary Table 13). Within these ANO3-ITPR1 clusters, the ANO3 and ITPR1 puncta were bimodally spaced between 10 nm and 30 nm of each other in both experimental systems (Figure 9l, Supplementary Table 13). This spacing is consistent with previously reported endoplasmic reticulum-plasma membrane (ER-PM) junction gap distances^41,42^. The bimodal distribution of our distance measurements could reflect the effect of cytoplasmic calcium fluxes, which have been previously shown to shrink the ER-PM junction gap from ∼25 nm to ∼10nm^43^.

Given the close spacing between ANO3 and ITPR1 both *in vitro* and *in vivo,* and previous reports of an ANO3 homolog, ANO1, physically interacting with the ITPR1 receptor^38^, we hypothesized that ANO3 could also physically interact with ITPR1. To address this question, we used an AlphaFold-predicted structure of ANO3 to identify intracellular domains within the protein that could possibly interact with the cytoplasmic-facing domains of ITPR1. In agreement with the previous ANO1 study^38^, we identified three main intracellular domains that are predicted to interact with cytoplasmic binding partners: the N-terminus, the first intracellular loop (ICL1), and the C-terminus. With this information, we used Alphafold3 to model the putative interaction between each of these three domains with the ITPR1 channel (Supplementary Table 14). While none of these domains were predicted to interact with ITPR1 with a high degree of confidence, the prediction confidence for ICL1 (chain ipTM=0.2) was notably higher than the prediction confidence for the other two cytoplasmic domains (C-terminus: chain ipTM=0.14; N-terminus: chain ipTM=0.13). Specifically, AlphaFold3 predicted that ANO3 ICL1 would interact directly with the cytoplasmic C-terminal domain of ITPR1, which composes the most cytoplasmic-facing portion of the ITPR1 calcium pore (Extended Data Figure 10a-b). This model would position the calcium pore of ITPR1 immediately adjacent to the putative allosteric calcium binding site of ANO3 (Figure 3m), which has been previously found to be critical in enhancing channel activity of the ANO1 and ANO6 homologs^44,45^.

To experimentally test the validity of this *in silico* prediction, we expressed constructs encoding the ICL1 sequence from ANO3 or ANO1 fused to green fluorescent protein (GFP), and a GFP-only control in iPSC-derived neurons (Figure 3n, Supplementary Table 14). The 55-amino acid ANO3 ICL1 domain, but not the ANO1 ICL1 domain, showed increased clustering with ITPR1 relative to the GFP control (Figure 3n). These findings support a model in which ANO3 physically interacts with ITPR1, involving interaction of the first intracellular loop of ANO3 with the cytoplasmic face of the calcium pore of ITPR1.

To test if ANO3-ITPR1 clusters are more active in exposing PtdSer than clusters containing only ANO3, as would be suggested by our model, we turned to expansion microscopy where we stained for exposed PtdSer via pre-fixation Annexin-V staining alongside staining for ANO3 and ITPR1 puncta. Nearest-neighbor analysis revealed that in human iPSC-derived neuron-astrocyte cocultures, both ANO3-ITPR1 and ANO3-only clusters significantly co-clustered with exposed PtdSer relative to background signal (Figure 3o, Supplementary Table 13). This suggests that these clusters likely affect PtdSer exposure in a spatially restricted fashion. We also found co-clustering of exposed PtdSer with independent ITPR1 clusters that was above background signal; however, given the heterozygous nature of the V5-tag knock in schema, our ANO3 labeling strategy would only label at most 50% of the ANO3 proteins (Figure 3o, Extended Data Figure 6a, Supplementary Table 13). Furthermore, we found that ANO3-ITPR1 clusters were significantly more likely than either independent ANO3 or independent ITPR1 clusters to co-cluster with exposed PtdSer both *in vitro* and *in vivo* (Figure 3p, Supplementary Table 13). This indicates a strong spatial correlation between physical ANO3-ITPR1 clustering events and increased local PtdSer exposure.

### ITPR1 activity drives synaptic phosphatidylserine exposure in an ANO3-dependent manner

Given these correlative results, we wanted to test if perturbing IP3 receptor activity would causally affect synaptic PtdSer exposure. We found that pharmacological inhibition of the IP3 receptor by Xestospongin C^46^ for three days was sufficient to substantially reduce synaptic PtdSer exposure in iPSC-derived neurons with no effect on synaptosome volume or somatic PtdSer exposure (Figure 4a,c; Extended Data Figure 5e). Conversely, three day-treatment with ARN11391, the ITPR1-specific positive allosteric modulator of channel activity^47^, increased synaptic PtdSer exposure without altering synaptosome volume or somatic PtdSer exposure (Figure 4b,d; Extended Data Figure 5f). This ITPR1-driven increase in synaptic PtdSer exposure was independent of caspase activity (Figure 4e), but required ANO3 (Figure 4f). These combinatorial perturbations subtly reduced synaptosome volume by ∼3% (Extended Data Figure 5g-h). These findings suggest that ITPR1 and ANO3 functionally associate to regulate synaptic PtdSer exposure.

**Figure 4.**
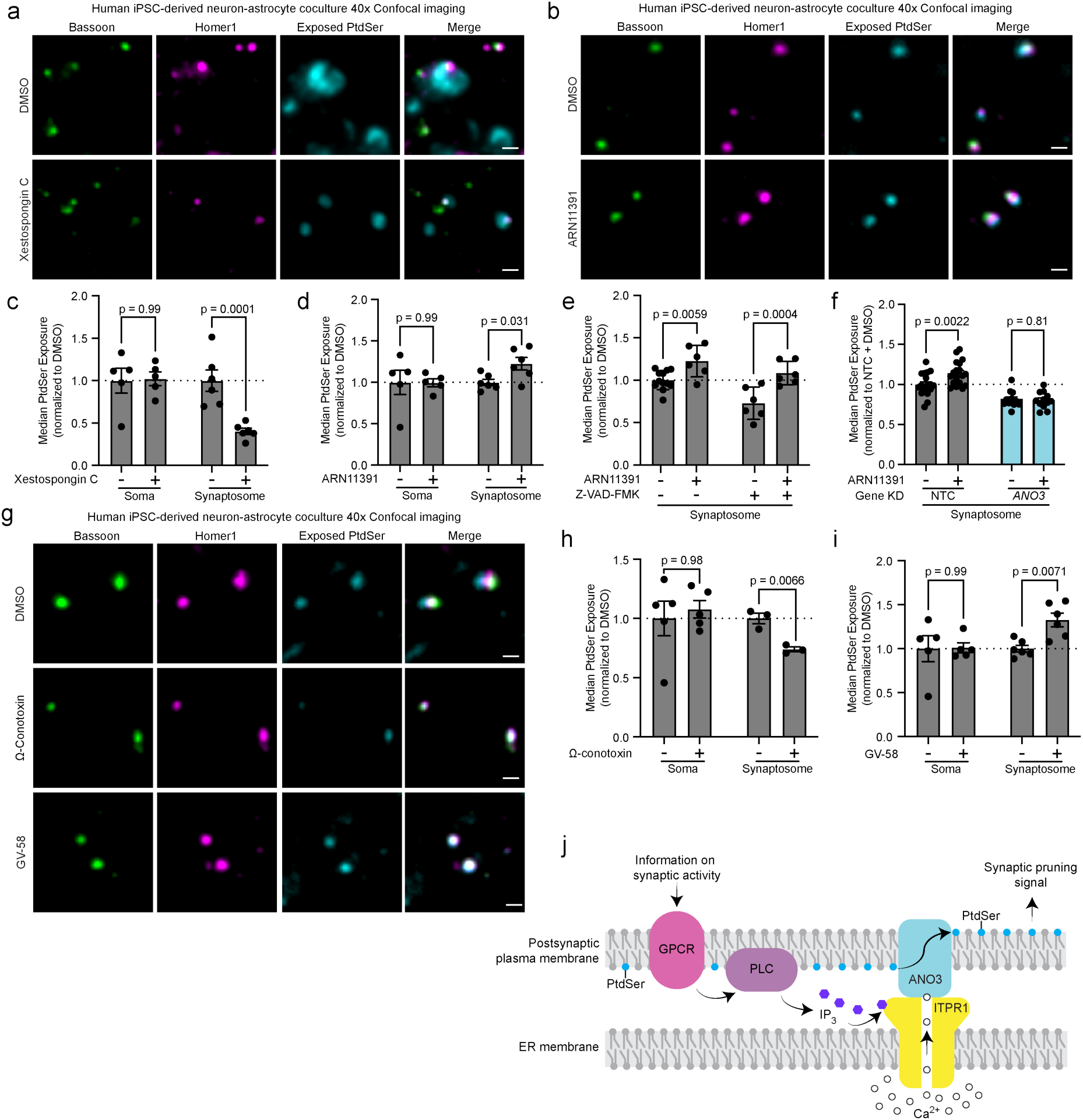
IP3 receptor activity modulates synaptic phosphatidylserine exposure in an ANO3 but caspase-independent manner. (a-b) Immunofluorescent staining for presynaptic (Bassoon) and postsynaptic (Homer1) markers alongside exposed phosphatidylserine (PtdSer) via pre-fixation Annexin V staining in human iPSC-derived neuron-astrocyte cocultures upon 72 hours of (a) IP3 receptor inhibition via 10 μM Xestospongin C treatment or (b) IP3R1 activation via 20 μM ARN11391 (Scale bar = 1 μm). (c-d) Flow cytometry reveals that (c) 72 hours of 10 μM Xestospongin C treatment selectively reduces synaptic PtdSer exposure, while (d) 72 hours of 20 μM ARN11391 selectively enhances synaptic PtdSer exposure (mean ± s.e.m., n=5-6 wells, p-values calculated by one-way unpaired followed by two-sided unpaired t-tests comparing against DMSO control with a Holm-Šídák correction for multiple comparisons applied). (e) 72-hour treatment of 20 μM ARN11391 still increases synaptic PtdSer exposure when simultaneously inhibiting caspase activity with 20 μM Z-VAD-FMK (mean ± s.e.m., n=6-12 wells, p-values calculated by two-way unpaired ANOVA with a follow-up Fisher’s LSD test, ARN11391 effect p-value < 0.0001, Z-VAD-FMK effect p-value = 0.0013, Interaction p-value = 0.267). (f) 72-hour treatment of 20 μM ARN11391 does not increase synaptic PtdSer exposure when simultaneously inhibiting ANO3 activity via CRISPRi-mediated ANO3 knockdown (mean ± s.e.m., n=12 wells, p-values calculated by two-way unpaired ANOVA with a follow-up Fisher’s LSD test, ANO3 knockdown effect p-value < 0.0001, ARN11391 p-value = 0.071, Interaction p-value = 0.031). (g) Immunofluorescent staining for presynaptic (Bassoon) and postsynaptic (Homer1) markers alongside exposed PtdSer via Annexin V (ANX) staining in human iPSC-derived neuron-astrocyte cocultures upon 72-hour treatment of 2 μM presynaptic voltage-gated calcium channel inhibitor 0-conotoxin or 72-hour treatment of 50 μM presynaptic voltage-gated calcium channel activator GV-58 suggests that synaptic activity can regulate synaptic PtdSer exposure (Scale bar = 1 μm). (h-i) Flow cytometry confirms that (h) 72 hour treatment of 2 μM presynaptic voltage-gated calcium channel inhibitor 0-conotoxin selectively reduces synaptic PtdSer exposure, while (i) 72 hour treatment of 50 μM presynaptic voltage-gated calcium channel activator GV-58 selectively enhances synaptic PtdSer exposure (mean ± s.e.m., n=3-6 wells, p-values calculated by one-way unpaired ANOVA followed by two-sided unpaired t-tests comparing against DMSO control with a Holm-Šídák correction for multiple comparisons applied). (j) Working model of molecular circuit by which synaptic activity is sensed by postsynaptic GPCRs to drive a postsynaptic IP3 response activating local IP3 receptors. This drives calcium release from the ER which directly activates colocalized ANO3 promoting local synaptic PtdSer exposure which can serve as a signal to glial cells to promote activity-dependent synaptic elimination.

### Synaptic activity bidirectionally regulates synaptic phosphatidylserine exposure

Postsynaptic IP3 receptors can release calcium in response to signaling from synaptic G protein-coupled receptors (GPCRs) that detect specific patterns of presynaptic neurotransmitter release^48^. Therefore, we hypothesized that presynaptic neurotransmitter release may also regulate synaptic PtdSer exposure. To test this hypothesis, we treated our human iPSC-derived neuron-astrocyte cocultures for 3 days with the presynaptic N-type voltage-gated calcium channel inhibitor 0-conotoxin^49^ or GV-58, a positive allosteric modulator of presynaptic N- and P/Q-type voltage-gated calcium channels^50^. 0-conotoxin inhibits synaptic vesicle release^49^ whereas GV-58 boosts synaptic vesicle release upon physiological presynaptic terminal depolarization^50^. We found that 0-conotoxin reduced synaptic PtdSer exposure whereas GV-58 increased synaptic PtdSer exposure within our human iPSC-derived neuron-astrocyte cocultures without affecting somatic PtdSer exposure (Figure 4g-i). The 0-conotoxin treatment did not affect synaptosome volume, while GV-58 slightly reduced synaptosome volume by ∼6% (Extended Data Figure 5g-h).

Based on our findings, we propose that presynaptic neurotransmitter release activates postsynaptic GPCRs to drive local ITPR1-mediated ER calcium efflux within the synapse. This calcium efflux activates local ANO3 at the postsynaptic membrane to drive synaptic PtdSer exposure, which ultimately signals glial cells to selectively eliminate synapses according to their synaptic activity patterns (Figure 4j).

## Discussion

### *In vivo* CRISPR screening platform for compartment-specific phenotypes

Here, we develop COMPASS-seq, a novel pooled screening platform for studying phenotypes of subcellular compartments. We apply COMPASS-seq here to uncover post-synaptic mechanisms controlling synapse numbers by using synapse-targeting nanobodies to localize guidecodes informing on genetic perturbations in the post-synaptic neuron. The platform could also be used to uncover presynaptic regulators, by using recently reported presynaptic traffickers^51^. Furthermore, the strategy makes it possible to investigate distinct types of synapses by using different Cre drivers to target neuronal subpopulations of interest.

Beyond the synapse, the screening approach employed in this study can be generalized to enable screens by modifying the trafficker to target guidecodes to other relevant subcellular structures, such as the myelin sheath or primary cilia, to systematically identify genetic regulators driving the formation and elimination of these structures. This screening platform can also be integrated with fluorescence-associated sorting screens to uncover genetic regulators of any subcellular phenotype that can be converted into a fluorescent readout. Lastly, this technology could also be integrated with spatial transcriptomics technology to enable pooled optical screening for regulators of neuronal morphology and subcellular transcriptomes.

### ANO3 regulates synaptic density and synaptic PtdSer exposure

Here, we identified ANO3 as the lone phospholipid scramblase knockdown of which increases synaptic density. ANO3 is expressed throughout neurons across the entire brain^52^ and is associated with various neurological conditions, including febrile seizures^8^ and DYT24 dystonia^9^. More recently, a meta-analysis identified mutations at the ANO3 locus as putative genetic drivers of dementia^7^. Our results suggest that causal association of ANO3 with these neurological diseases may be mediated by the mechanistic role of ANO3 in regulating synaptic density.

Based on overexpression studies in immortalized cell lines, ANO3, like several other anoctamins, can act as a calcium-dependent phospholipid scramblase^28^. However, it was previously unknown where ANO3 functions in a physiological context. We found that synaptic ANO3 tends to localize to the postsynaptic membrane, compatible with the role we uncover for ANO3 in controlling synaptic PtdSer exposure. However, a substantial fraction of ANO3 was localized extrasynaptically. It is possible that extrasynaptic ANO3 also contributes to its effect on synaptic density, possibly through its previously described role in regulating neuronal excitability via a physical interaction with sodium-activated potassium channels^53^.

### ITPR1 activity drives synaptic PtdSer exposure in an ANO3-dependent manner

ANO3 has been previously shown to be activated by calcium^28^. In search of a postsynaptic calcium source that may physiologically drive ANO3 activity at the postsynaptic membrane, we identified ITPR1, an IP_3_-activated endoplasmic reticulum calcium channel, as a driver of ANO3 activity. ITPR1 has previously been identified as a key regulator of mGluR-dependent synaptic long-term depression (LTD)^39,40^. Here, we associate ITPR1 signaling with a novel mechanism through which it may promote glia-dependent synapse loss. Several groups have previously hypothesized that there could be a mechanistic link between synaptic long-term depression and glia-mediated synapse loss^15,17,18^. Based on our findings in this study, we propose that ITPR1 could serve as one of the mechanistic nodes via which both synaptic LTD and glia-mediated synapse loss could be coregulated.

ITPR1 is activated by IP_3_, which is generally produced in response to Gq-GPCR signaling. A candidate Gq-GPCR activating the ITPR1-ANO3 pathway is mGluR5. Postsynaptic mGluR5 can be aberrantly activated by amyloid-beta oligomers binding to the prion protein^54^, and mGluR5 signaling can promote complement tagging of synapses and downstream glia-mediated synapse loss in the context of Alzheimer’s disease models^55,56^. Aberrant activation of mGluR5 could engage the ITPR1-ANO3 pathway discovered here to enhance PtdSer exposure, which forms an extracellular binding site for complement proteins promoting glia-mediated synapse loss^22^.

ANO3 scramblase activity may be further regulated by other signals. While biophysical studies of ANO3 regulation are limited, the homologous proteins ANO1 and ANO6 have previously been shown to require PIP_2_ at the plasma membrane to maintain sustained activity^57,58^. If ANO3 activity is similarly regulated by plasma membrane PIP_2_, this could form the basis of a homeostatic feedback loop where PIP_2_ cleavage into IP_3_ and diacylglycerol can produce a self-limiting amount of synaptic PtdSer exposure via the ITPR1-ANO3 interaction discovered here. ANO1 activity has also been found to be dependent on membrane potential^59^, which may also further regulate ANO3 function.

### Role for synaptic activity in regulating glia-mediated synapse loss

We also found that increased synaptic neurotransmitter release enhanced synaptic PtdSer exposure. This is consistent with prior reports that hyperactive synapses preferentially expose phosphatidylserine^6,25^, and that induction of presynaptic neuronal activity can promote both synaptic PtdSer exposure and glia-mediated synapse loss^6,25^. Other studies have found that inactive synapses are preferentially engulfed by glial cells^5,18^. It seems likely that several orthogonal activity-dependent mechanisms regulating exposure of “Eat-me” and “Don’t-eat-me” signals on synapses may be converging to control the relationship between synaptic activity and synapse loss in different contexts. Amongst these mechanisms are previous reports on the role of activity in regulating phospholipid flippase cochaperone TMEM30A expression^60^ and CD47 localization^61^. Future work is needed to precisely understand how the ANO3-ITPR1 mechanism described here interacts with these other known mechanisms to ultimately regulate glia-mediated synapse exposure and neuronal circuit changes both in physiological contexts as well as in the context of neurological disease.

## Methods

### Plasmid generation

pTS026 was generated using pAP215^33^ as the starting backbone. We subcloned the mU6-sgRNA-Cre-invertible handle region from its initial position on the positive strand upstream of the EF1a promoter onto the negative strand downstream of the Ef1a promoter. Next, we replaced the Ef1a promoter with the hSYN promoter from pENN.AAV.hSyn.HI.eGFP-Cre.WPRE.SV40 (Addgene #105540, a gift from James M. Wilson). Then, we replaced the NLS-mTagBFP2 coding sequence and invertible handle region with a Cre-on flip-excision system using a Gene Fragment (Twist Biosciences) based on a previously reported sequence^62^. Next, we subcloned the PSD95.FingR.eGFP.CCR5TC coding sequence from Lenti-CamKII-PSD95.FingR-eGFP-CCR5TC^63^ (Addgene #126218, a gift from Xue Han) in an inverted orientation within the flip-excision cassette and the zinc-finger binding domain sequence from that same plasmid upstream of the hSYN promoter. Then, we replaced the eGFP sequence with a poly-GA linker- nλ – poly GA linker–mScarlet-I– poly GA linker sequence using a Gene Fragment (Twist Biosciences) based on the mScarlet-I sequence from pJC49^64^ and the nλ domain sequence from IDP424^65^ (Addgene #87018, a gift from Anthony Zador). Finally, we added the 4xBoxB sequence from jk100 pSinEGdsp (Addgene #79785, a gift from Anthony Zador) in an inverted orientation within the flip excision cassette downstream of the PSD95-FingR coding sequence stop codon. To enhance library cloning efficiency, an optimized protospacer sequence (GACCCGACTTTAATTAAGGA) was cloned into the sgRNA scaffold.

To prepare pKS141, the mScarlet-I sequence in pTS026 was replaced with a Halotag sequence from pAJS212^66^. pKS145 was generated by removing the 4xBoxB domain from pKS141 via restriction digestion and religation.

For knockdown of target genes within human iPSC-derived systems, individual sgRNAs were cloned into the pMK1334 backbone as previously described^67^. Protospacer sequences used in this manuscript are listed in Supplementary Table 7.

GFP fusion constructs were generated from a pJC49 backbone. Superfolder GFP was subcloned into pJC49^64^ via Gibson Assembly of a Gene Fragment (Twist Bioscience) to produce pKS148. Next, the first intracellular loop of ANO1 and the first intracellular loop of ANO3 were subcloned from a Gene Fragment (Twist Bioscience) to fully prepare pKS151and pKS236, respectively. Peptide and DNA sequences for the first intracellular loop of ANO1 and ANO3 are detailed in Supplementary Table 14.

### Flippase and scramblase library cloning

A custom library encoding a 275 element sgRNA library composed of 150 non-targeting control sgRNAs and 125 sgRNAs targeting all 28 mouse flippase and scramblase genes was purchased from Twist Bioscience. Protospacer sequences for this library are listed in Supplementary Table 7. This library was cloned into pTS026 containing the optimal sgRNA protospacer for efficient cloning using the optimized library cloning procedure from Heo et al. (2024)^68^ with the following minor modifications. In addition to Bpu1102I (Thermo Scientific, Cat# FD0094) and BstXI (Thermo Scientific, Cat# FD1024), pTS026 was additionally digested using PacI (Thermo Scientific, Cat# FD2204) to degrade the initial protospacer sequence contained within pTS026. The linearized vector was purified using a DNA Cleanup Kit (NEB, Cat# T1130L).

Ligation was performed to generate library of protospacers inserted into linearized pTS026. Each 20 μL ligation reaction contained 100 ng of linearized vector DNA, 0.68 ng of protospacer library insert DNA, and 2 μL T4 DNA Ligase (NEB, Cat # M0202S). Ligation products were subsequently purified using a DNA Cleanup Kit. To ensure adequate library representation, test transformations were performed to evaluate ligation and transformation efficiencies. Finally, library coverage and diversity were validated via Next-Generation Sequencing (NGS) on the Illumina NextSeq 2000 platform.

### AAV packaging

AAV was packaged, purified, and titered as previously described^33^. Briefly, HEK293T cells (ATCC, CRL-3216) were transfected with pAdDeltaF6 (Addgene #112867, a gift from James M. Wilson), pUCmini-iCAP-PHP.eB^69^(Addgene #103005, a gift from Viviana Gradinaru), an AAV transfer plasmid, and polyethylenimine (PEI, Polysciences, Cat #23966), all diluted in 5 mL of Opti-MEM (Gibco, Cat #31985062). The mixture was incubated at room temperature for 10 minutes and then added dropwise onto HEK293T cells. 72 hours after transfection, cells and media were triturated, collected, and chloroform (Sigma-Aldrich, Cat #C2432) extracted. Viral particles were precipitated from the aqueous phase overnight before being pelleted by centrifugation and resuspended in 1 mL of 50 mM HEPES buffer (Sigma-Aldrich, Cat #H3375). supplemented with 3mM of MgCl_2_. This solution was treated with RNase (Thermo Scientific, Cat #EN0531) and DNase (NEB, Cat #M0303S) followed by a second round of chloroform extraction. The aqueous phase was then passed through a 0.5mL Amicon Ultra Centrifugal Filter with a 100 kDa cutoff (Millipore, Cat #UFC510024). 1X PBS (Sigma-Aldrich, Cat# D8537) was then passed through the same filter to perform a buffer exchange prior to elution. Titering was completed by quantitative RT-PCR using primers ITR-FWD and ITR-REV targeting the ITR of the AAV plasmid. Primer sequences are listed in Supplementary Table 3.

### Lentivirus packaging

Lentivirus was packaged and stored as previously described^67^. Briefly, HEK293T cells were transfected with 3^rd^ generation lentiviral packaging plasmids, a lentiviral transfer plasmid, and PEI all diluted in 200 μL of Optimem was prepared. The mixture was incubated at room temperature for 10 minutes and then added dropwise onto HEK293T cells. 48 hours after transfection, supernatant was taken from each well and filtered. Filtered supernatant was supplemented with lentivirus precipitation solution (ALSTEM, Cat #VC100) and incubated at 4°C for three days to precipitate viral particles. Precipitated viral particles were pelleted and resuspended in 500 μL 1X PBS prior to being used for downstream applications.

### Mouse husbandry

All mice were maintained according to the National Institutes of Health guidelines. All procedures were approved by the University of California, San Francisco Institutional Animal Care and Use Committee. Mice were housed under 22–25 °C and 50–60% humidity and a 12 hour light/12 hour dark cycle. Food and water were provided ad libitum. Mice were randomly assigned to the experimental groups at the time of experiment, and both male and female mice were used. Experimental details of the mice used in each figure are listed in Supplementary Table 1. The mice used in this study include: LSL-dCas9-KRAB (LSL-CRISPRi) mice (B6;129S6-Gt(ROSA)26Sortm2(CAG-cas9*/ZNF10*)Gers/J, RRID: IMSR_JAX:033066) and the newly developed *Ano3* knock-in mouse described below.

### *Ano3* knock-in mouse line development and validation

An *Ano3* knock-in mouse line was generated using a CRISPR/Cas9-mediated genome editing strategy to introduce a C-terminal epitope tag immediately upstream of the endogenous stop codon of *Ano3*. The donor construct was generated using the pFETCH donor backbone (Addgene #63934, a gift from Eric Mendenhall and Richard M. Myers) and contained a 5-amino-acid linker followed by a FLAG-GFP11-FLAG-GFP11-FLAG.

The inserted DNA sequence was:

GGTGGCGGCAAATTCGATTACAAGGATGACGACGATAAGGGTCGTGACCACATGGTCCTTCATGAGTATGTAAATGCTGCTGGGATTACAGGCGATTATAAGGACGACG ATGATAAAGGTCGAGACCATATGGTTCTCCACGAATACGTAAACGCAGCAGGCATCACTGGAGACTACAAAGACGATGACGACAAGTGA

The knock-in cassette was inserted immediately before the endogenous stop codon to preserve endogenous *Ano3* regulatory and coding sequences while enabling C-terminal tagging of *Ano3*. Founder mice carrying the correctly targeted allele were identified by genotyping using the following primer pairs: mouse_A3KI_F1, mouse_A3KI_R1 or mouse_A3KI_F2, mouse_A3KI_R2 (Extended Data Figure 6b). Primer sequences are listed in Supplementary Table 3. Genotyping PCR reactions consisted of 1x GoTaq Master Mix (Promega, Cat #M7123) supplemented with 500 nM of the FWD primer, 500 nM of the REV primer, and 2 μL of 1:100 diluted DNA collected from a tissue sample. The thermal cycling conditions consisted of an initial 2 min at 94°C, followed by 35 cycles of 30 sec at 94°C, 60 sec at 60°C, and 45 sec at 72°C, followed by a final 2 min extension at 72°C. The knock-in allele was sequenced to confirm successful in-frame insertion of the tag.

### Neonatal intracerebroventricular injections

Injections were performed as previously described^33^. Briefly, neonatal mouse pups were cryoanesthetized and the scalp was swabbed with an alcohol wipe. AAVs were diluted to desired titer in 1X PBS to a volume of 2 μL, with 0.1% Trypan Blue (Invitrogen, Cat #T10282) to visualize injection spread into ventricles at time of injection. Virus solution was slowly injected into the ventricle unilaterally or bilaterally by hand, and the needle was left in place for up to 30 seconds to prevent backflow of solution. Following the injection, the neonate was placed on a warming pad until signs of recovery were observed and then returned to parent cage. All injection information for specific mice is listed in Supplementary Table 1.

### Adult Annexin-V intracerebroventricular injections

Adult mice were anesthetized using 2-3% isoflurane (Kent Scientific, J124013) + oxygen mixture and injected with extended-release buprenorphine (Ethiqa-XR, 86084-10-30), carprofen (Rimadyl, 11831), and lidocaine (Vedco, 50989-0417-15) at dosages outlined by UCSF IACUC and LARC guidelines for pain management in rodents. Mice were subsequently positioned within the stereotactic frame (David Kopf, Model 940) and the scalp fur was trimmed and swabbed with betadine (Dynarex, 1415) and alcohol (First Aid Only, H305-001) to sterilize the surgical field. After confirming anesthesia depth, an incision was made on the scalp to expose the skull, and the head was aligned relative to the frame using bregma and lambda. Once appropriately positioned, a drill attachment (David Kopf, Model 1474) was used to drill a hole in the skull at −0.46 AP, 1.0 ML, −2.3 DV to target the ventricle. The drill attachment was swapped for a microinjector (World Precision Instruments, UMP3 Model) equipped with a syringe (Hamilton, 7635-01) loaded with a pulled glass micropipette and 2 uL of Annexin-FITC dye (Invitrogen, Cat #A13199) was slowly injected into the ventricle unilaterally. The microinjector needle was left in place for 2 minutes to prevent backflow of solution. Following the injection, the initial incision was closed with Vetbond (Patterson Veterinary Supply, 07-805-5031) and removed from the stereotactic frame. Mice were placed on a warming pad (Kent Scientific HTP-1500) for recovery and then returned to their home cage. Mouse brains were collected the next day for downstream applications. All injection information for specific mice is listed in Supplementary Table 1.

### Human iPSC culture

IPSC culture was performed as previously described^67^. Briefly, WTC11 background human iPSCs were cultured in Stemflex media (Gibco, Cat #A3349401) on BioLite Cell Culture Treated Dishes (Thermo Scientific) coated with Growth Factor Reduced, Phenol Red-Free, LDEV-Free Matrigel Basement Membrane Matrix (Matrigel, Corning, Cat #356231) diluted 1:100 in Knockout DMEM (Gibco, Cat #10829-018). Stemflex media was replaced every other day or every day once 50% confluent. When 80-90% confluent, cells were passaged, which entailed the following: aspirating media, washing with 1X PBS, incubating with StemPro Accutase Cell Dissociation Reagent (Gibco, Cat #A1110501) at 37°C for 10-15 minutes, diluting with 1X PBS, collecting dissociated cells in conical tubes, centrifuging at 220xg for five minutes to pellet the cells, resuspending the pellet in Stemflex media supplemented with 10 nM Y-27632 dihydrochloride (ROCK inhibitor, Tocris, Cat #125410) and plating onto Matrigel-coated plates. Studies with human iPSCs at UCSF were approved by the Human Gamete, Embryo and Stem Cell Research (GESCR) Committee. Informed consent was obtained from the human subjects when the WTC11 line was originally derived^70^.

### Astrocyte differentiation

Astrocyte differentiation was performed as previously described^71^. Briefly, CD133+/CD271− sorted NPCs generated from WTC11 background human iPSCs with stably integrated CRISPRi machinery and dox-inducible NFIA-SOX9 were plated at 7,500–15,000 cells per cm^2^ in NPC media^71^ onto a 10-cm or 15-cm dish coated with Matrigel diluted at 1:200 in Knockout DMEM. The next day, media was changed to astrocyte media (ScienCell, Cat #1801) supplemented with 2 μg/mL doxycycline (Sigma-Aldrich, Cat #D9891) to initiate iAstrocyte differentiation. Media was fully changed every other day, with doxycycline maintained at 2 μg/mL throughout the differentiation process. When the differentiating NPCs reached confluency every 3–4 days, the culture was dissociated with StemPro Accutase cell dissociation reagent and replated onto new Matrigel coated dishes at a plating density of 10,000–20,000 cells per cm^2^. Expansion of the cultures was continued until day 20 of differentiation, yielding iPSC-derived astrocytes, which were used for downstream functional assays.

### Neuron-astrocyte coculture plating and maintenance

To prepare cocultures, WTC11 background human iPSCs were passaged once before beginning the differentiation protocol. Neuronal pre-differentiation was performed as previously described^67^. Briefly, cells were grown for three days in predifferentiation media composed of Knockout DMEM/F12 (Gibco, Cat #12660-012), 1X non-essential amino acids (Gibco, Cat #11140-050), 1X N-2 Supplement (Gibco, Cat #17502048), 10 ng/mL BDNF (Peprotech, Cat #450-02), 1 μg/mL mouse laminin (Thermo Scientific, Cat #23017015), 10 ng/mL NT-3 (Peprotech cat #450-03), and 2 μg/mL doxycycline. For the first 24 hours, 10 nM ROCK inhibitor was added to the predifferentiation media after which the media is replaced with fresh predifferentiation media not containing ROCK inhibitor.

Cocultures were plated onto plates coated with Poly-D-Lysine (PDL) (Sigma-Aldrich, Cat #P7405) and mouse laminin using the following coating protocol. On day –3 of differentiation, plates were coated with 1 mg/mL PDL resuspended in 1X borate buffer (Thermo Scientific, Cat #28341) and stored at 37°C overnight. On day −2, the plates were washed four times with cell culture grade H_2_O (Corning, Cat #25-055-CVC) and left to air dry overnight. On day –1, the plates were coated with 10 μg/mL mouse laminin diluted in 1X PBS with calcium and magnesium added (Corning, Cat #20-030-CV) Plates were incubated at 37°C overnight.

On day 0, predifferentiated neurons were replated with D20-D21 astrocytes at a 2:1 neuron: astrocyte ratio at a total cell density of 700,000 cells per cm^2^. Cells were plated in a 1:1 mix of neuronal media with astrocyte media. Neuronal media was prepared from BrainPhys (STEMCell Tech, Cat #05791) supplemented with 0.5X N-2 supplement, 0.5X B-27 supplement minus Vitamin A (Gibco, Cat #12587010), 10 ng/mL BDNF, 1 μg/mL mouse laminin, and 10 ng/mL NT-3. Immediately after replating, 2 μg/mL doxycycline was added to coculture media. On day 3 after replating, plating media was aspirated and replaced with fresh neuronal media supplemented with 2 μg/mL doxycycline. After this media change, the cells were fed every 48 hours in a half media change of fresh neuronal media without doxycycline supplementation.

### CRISPR editing of iPSCs

Human iPSCs were edited following the previously described protocol^72^. A single-stranded oligodeoxynucleotide encoding a V5 tag and a flexible linker sequence with homology arms targeting insertion of the tag immediately upstream of the ANO3 start codon to preserve the regulatory regions within the 3’ UTR of ANO3. Genomic DNA was targeted with the following modified SpCas9 gRNA: ACUGAAUGGAGCCUGAAUGG (Synthego). The single-stranded oligodeoxynucleotide was synthesized (Integrated DNA Technology) with the following sequence:

AGGGGAGCAAGATTCCAACTGACCTTTTTGCTGTTTAAAGGACTGGATTGATCCGGAGTGATGCACCATGCCAGCTCCGGCTCCAGCTCCAGCGCCAGCTCCGGCTCCCGTAGAATCGAGACCGAGGAGAGGGTTAGGGATAGGCTTACCCATTTTCACTCTGCGCGTCCCAGCCGGCTTAGGGAGCTGCCCGAGAGGGA

Individual colonies were genotyped for successful CRISPR editing with human_A3KI_FWD and human_A3KI_Rev genotyping primers (Extended Data Figure 6a). Primer sequences are listed in Supplementary Table 3. Genotyping PCR reactions were prepared as 1x GoTaq Master Mix supplemented with 200 nM of the FWD primer, 200 nM of the REV primer, and 1 μL of DNA collected from a cell lysate sample. The thermal cycling conditions consisted of an initial 2 min at 94°C, followed by 36 cycles of 60 sec at 94°C, 60 sec at 63°C, and 60 sec at 72°C followed by a final 10 min extension at 72°C. The knock-in allele was sequenced to confirm successful in-frame insertion of the tag.

### Drug treatments

On Day 18 post-plating of the coculture, cell culture media was fully replaced with fresh neuronal media supplemented with specified drug treatment or vehicle control at the working concentrations listed in Supplementary Table 10. On Day 20 post-plating of the coculture, a half-media change was performed with fresh neuronal media supplemented with either drug treatment or vehicle control at the working concentration. On Day 21 post-plating of the coculture, samples were collected for downstream analysis.

### Multielectrode array electrophysiology

Cocultures were plated on 24-well MEA plates (Axion, Cat #M384-tMEA-24W) coated with PDL and laminin as previously described. 110,000 predifferentiated neurons were mixed with 55,000 D20-21 astrocytes in a 20 μL media droplet placed on the central electrode region. One hour after plating, a 1 mL mix of 1:1 neuronal media and astrocyte media was added to each well. Media was changed as described above on Day 3 post plating and every other day after that. From Day 7 onwards, after changing the media, the plate was placed in the MEA system and allowed to acclimate at 37°C for 15-30 minutes. Following acclimation, 12 consecutive 5-minute recordings were collected from each MEA plate. When applicable, drug treatments were applied on Day 18 post-plating and treatments were not readministered during the following media changes.

Following the conclusion of the experiment, spike analysis was performed on the activity recordings using a static threshold of 10 μV. The average spike rate and the area under the normalized cross correlogram (AUNCC) were calculated for each well on each day of recording using the AxIS Navigator v2.0.4 software package along with Axion Biosystems Neural Metrics Tool v2.5.1. Data across the twelve 5-minute recordings were pooled together using a custom script to quantify the average spike rate and AUNCC for each well for each day of recording.

### Neuron-astrocyte coculture fixation

When visualizing exposed PtdSer for immunofluorescent imaging, prefixation annexin staining was performed by diluting Annexin-647 (Invitrogen, Cat #A23204) 1:200 in fresh neuronal media and incubating at 37°C for 1 hour. All buffers for subsequent wash and incubation steps were prepared in 1X PBS with calcium and magnesium added. After incubation, cells were washed once and then subsequently fixed with 4% paraformaldehyde (PFA, Electron Microscopy Sciences, Cat #50-980-495) for 10-15 minutes at room temperature. Following fixation, cells were washed 3 times. For expansion microscopy, cells were post-fixed in 0.7% PFA /1% acrylamide (AAm, Supelco Cat #01697) solution for 6 hours at 37°C followed by 2 washes with wash buffer. Fixed cells were stored in wash buffer at 4°C until use.

### Neuron-astrocyte coculture immunostaining

Blocking solution was prepared from 1X PBS with calcium supplemented with 5% normal goat serum (Gibco, Cat #16210072), and 0.01% Triton X-100 (Sigma-Aldrich, Cat #T9284). Cells were incubated with blocking solution for 2 hours at room temperature with agitation. Cells were incubated with blocking solution supplemented with primary antibodies for 48 hours at 4°C with agitation. After incubation, three washes were performed with agitation. Secondary antibodies were diluted in blocking solution at a 1:1000 dilution. Cells were incubated in blocking solution supplemented with secondary antibodies for 2 hours. Finally, three washes were performed with agitation. Sections were then mounted using either Prolong Gold (Life Technologies, Cat #P36930) for confocal imaging or Prolong Glass antifade mountant (Invitrogen, Cat #P36980) for STED super-resolution imaging and stored in darkness at 4°C until imaging. All antibodies used and associated concentrations are listed in Supplementary Table 2.

### Perfusion, fixation and sectioning of mouse brain tissue

Animals were transcardially perfused following UCSF IACUC guidelines. Animals were deeply anesthetized using a mixture of oxygen and isoflurane (3% flow rate), and lack of bipedal rear paw pinch reflex was used to confirm anesthesia depth. An incision was made across the abdomen to expose the thoracic cavity, and the diaphragm was ruptured. A needle with tubing attached (Fisher Scientific, Cat #22-258091) was then placed into the left ventricle of the heart, the right atrium cut, and 10-20 mLs of 1X PBS pumped through the vascular system using a syringe pump (rate of 2 mL/min, World Precision Instruments, Cat #AL-4000) until all blood was removed from the system. 4% PFA in 1X PBS was then pumped through the vascular system at the same rate and volume, until signs of fixation were observed. The brain was isolated from the skull, post-fixed overnight in 4% PFA in 1X PBS. The following day, the solution was removed and exchanged with 30% sucrose (Fisher Scientific, Cat #S5-500) in 1X PBS for cryopreservation, which brains were stored in until sectioning.

Brains were frozen in blocks of OCT (Fisher Scientific, Cat #23730571) media and sectioned using a Leica Microsystems CM3050S Cryostat to a thickness of 16-30um. Sections were stored free-floating in 24-well plates in cryopreservation media (50% ethylene glycol (Sigma-Aldrich, 102466) + 1% PVP (Sigma-Aldrich, PVP40-50G) in 1X PBS at 4°C.

For expansion microscopy, mice were perfused as previously described^37^ first with ice-cold 1X PBS and then with an ice-cold mixture of 4% PFA + 20% AAm. Brains were isolated and fixed overnight in 4% PFA + 20% AAm at 4°C, then washed and stored in 1X PBS until sectioning. Brains were sectioned to a thickness of 70 μm using a vibratome. Sections were stored in 1X PBS at 4°C until use.

### Mouse brain tissue immunostaining

Free-floating brain sections were mounted onto sample slides (Fisher Scientific, Cat #1255015). Sections were first washed 3 times with 1X PBS, then 3 times with PBST (1X PBS + 0.1% Triton-X 100). Sections were then blocked at room temperature for 1 hour using blocking solution (1% bovine serine albumin (Fisher Scientific, Cat #BP9700100) + 0.3% Triton-X 100 + 10% Normal Goat Serum (NGS) in 1X PBS). Primary antibodies, diluted in 1% BSA + 0.3% Triton-X 100 + 5% NGS in 1X PBS to a concentration of 1:500, were then applied to sections and incubated for 48 hours at 4°C. Sections were then washed 3 times with 1X PBS and 3 times with PBST. Secondaries were diluted in 1% BSA + 0.3% Triton-X 100 + 5% NGS in 1X PBS to a concentration of 1:1000 and incubated for 2-3 hours at room temperature in darkness. The nuclear marker Hoechst 33342 (Invitrogen, Cat #62249) was diluted to 1:1000 in 1X PBS and sections were incubated with this solution for 5 minutes at room temperature in darkness. Sections were then washed 3 times with PBST and 3 times with 1X PBS to remove excess antibody.

To simultaneously stain multiple rabbit-host antibodies as done in (Figure 3e-f, Extended Data Figure 7e,g), anti-GFP antibody (Invitrogen, Cat #PA5-109258) were conjugated to Biotin using the FlexABLE conjugation kit (Proteintech, Cat #KFA007). To mitigate non-specific background staining, sections were additionally blocked using Streptavidin/Biotin blocking solution (Invitrogen, Cat #37628) prior to staining. Following blocking, the conjugated antibody solution was then diluted into 1% BSA + 0.3% Triton-X 100 + 5% NGS in 1X PBS to a final concentration of 1:200 and sections were incubated for 24 hours at 4°C, and then washed 3 times with 1X PBS to remove excess antibody. Next, Alexa Fluor 488-conjugated streptavidin (Invitrogen, Cat #S32354) was diluted in 1% BSA + 0.3% Triton-X 100 + 5% NGS in 1X PBS to a concentration of 1:1000 and incubated for 2-3 hours at room temperature in darkness. Sections were then washed 3 times with PBST and 3 times with 1X PBS to remove excess conjugated streptavidin.

Sections were then mounted using either Prolong Gold for confocal imaging or Prolong Glass antifade mountant for STED super-resolution imaging and stored in darkness at 4°C until imaging. All antibodies used and associated concentrations are listed in Supplementary Table 2.

### Expansion microscopy sample processing

Medial hippocampal sections from perfused mice and cell samples grown on 25-mm coverslips (Neuvitro, Cat #GG-25-1.5h-pre) were selected and prepared for expansion microscopy as previously described^37^. Briefly, samples were first incubated in inactivated first gelling solution (10% AAm, 0.1% DHEBA (Sigma-Aldrich, Cat #294381), 19% sodium acrylate (Pfaltz and Bauer, Cat #101181-226) in 1X PBS) for 30 minutes on ice on an orbital shaker (IBI Scientific, Cat #BDRAA115S), and then incubated in activated first gelling solution (inactivated first gelling solution supplemented with 0.05% APS (Sigma-Aldrich, Cat #A3678) and 0.05% TEMED (Thermo Scientific, Cat #17919) for 15 minutes on ice on an orbital shaker. Gels were then transferred to a gelation chamber, placed into the humidified gas exchange chamber, and perfused with nitrogen gas for 10 minutes, before being incubated for 2 hours at 37°C. Biopsy punches (Sklar, Cat #96-0192) were used to extract several 2 mm diameter gels from each sample. For tissue samples., punches were aimed at the CA1 region of hippocampus. Gels were then transferred to 2 mL tubes containing 1 mL of denaturation buffer (200 mM NaCl (J.T. Baker, Cat #3624-01), 50 mM Tris (Genesee Scientific, Cat #18-190), 200 mM SDS (Invitrogen, Cat #AM9820) in ultrapure water) and incubated on a thermomixer at 75°C for either 4 hours for tissue-containing gels or for 1 hour for cell culture-containing gels. After denaturation, gels were washed 3 times with 1X PBS for 10 minutes each, 2 times with ultrapure water for 30 minutes each, and once with ultrapure water overnight.

Next, gels were incubated in freshly prepared 2^nd^ gelling solution (10% AAm, 0.05% DHEBA, 0.05% APS, 0.05% TEMED in ultrapure water) three times for 20 minutes at room temperature. Gels were then transferred to a gelation chamber, placed into the humidified gas exchange chamber, and perfused with nitrogen gas for 10 minutes, before being incubated for 2 hours at 37°C. Gels were placed into 1X PBS overnight at room temperature.

Next, gels were incubated in freshly prepared 3^rd^ gelling solution (10% AAm, 0.1% BIS (Alfa Aesar, Cat #J66710-14), 19% sodium acrylate, 0.05% APS, 0.05% TEMED in 1X PBS) three times for 15 minutes each on ice. Gels were then transferred to a gelation chamber, placed into the humidified gas exchange chamber, and perfused with nitrogen gas for 10 minutes, before being incubated for 2 hours at 37°C. Gels were then incubated in 200mM sodium hydroxide (NaOH; Sigma-Aldrich Cat #S8045) for 1 hour at room temperature, and then washed 3 times in 1X PBS for 30 minutes each. The regions of interest were then excised from the gels using a razor blade. Next, gel samples were washed with PBS-Tween (0.1% (w/v) Tween-20 (Sigma-Aldrich, Cat #P2287) in 1X PBS) once and incubated with antibody blocking buffer supplemented with primary antibodies diluted 1:250 for 48-72 hours on an orbital shaker at room temperature. Samples were washed three times with PBS-Tween for 20 minutes each and then incubated with antibody blocking buffer supplemented with secondary antibodies diluted 1:250 for ∼72 hours on an orbital shaker at room temperature protected from light. All antibodies used and associated concentrations are listed in Supplementary Table 2.

For experiments requiring pan-staining, samples were incubated with 20ug/mL STAR 635P-NHS ester (Abberior Star, Cat #ST635P-0002-1MG) diluted in 100 mM sodium bicarbonate (Sigma-Aldrich, Cat #S5761) for 2 hours. For experiments requiring SYTOX green staining, gel samples were incubated with SYTOX green (Invitrogen, Cat #S7020) diluted 1:3000 in HBSS buffer (Gibco, Cat #14170112) for 30 minutes on an orbital shaker at room temperature. Gel samples were then washed three times with PBS-Tween for 20 minutes each on an orbital shaker at room temperature. Finally, gel samples were washed twice in ultrapure water for 1 hour per wash followed by a final wash in ultrapure water overnight.

Next, gels were sliced and mounted on glass-bottom MatTek dishes (MatTek Life Sciences, Cat #P50G-1.5-30-F). A #1.5 40 x 24 mm coverslip (Epredia, Cat #152440) was placed over the gels, and silicone glue (Henry Schein, Cat #1026526) was used to seal the samples. Mounted samples were then stored in the dark until imaging. All imaging of expanded samples was performed within 72 hours of mounting.

### Neuron-astrocyte coculture high content confocal microscopy

High-content imaging of fixed neuron-astrocyte coculture samples was performed using an ImageXpress Confocal HT.ai Imaging System (Molecular Devices) running MetaXpress v6.7.1.157 equipped with an LDI laser combiner (89North) and a Zyla 4.2Plus camera (Andor). Images were acquired using a Plan Apo LWD 40x/1.15NA WI objective in confocal mode (60 μm pinhole) with excitation lasers of 475 nm, 555 nm, and 730 nm wavelengths and collected using pre-set filters for these channels. 4-9 images were collected per well, and 3-6 wells were imaged per condition. No deconvolution was used to process any confocal images. For representative images within this manuscript, bilinear interpolation was used to adjust the resolution to >300 dpi.

### High-resolution confocal microscopy

Imaging of *in vivo* fixed samples was performed using a spinning disk confocal microscope equipped with a Yokogawa CSU-W1 SoRa module set on a DMI8 inverted microscope (Leica Microsystems). 488 nm, 561 nm and 639 nm lasers (Versalase, Vortran) were used for excitation and emission was collected using appropriate bandpass filters on a Teledyne Kinetix sCMOS camera. Multichannel images were acquired sequentially to minimize spectral crosstalk. Acquisition was done using the custom Micro-Manager Studio v.2.0.3 software system^73^. Images of standard slide-mounted brain tissue samples were acquired using an HC PL APO 100x/1.47 OIL objective, and from each sample, images of 4-6 regions of the hippocampus including the dentate gyrus, the CA1 and the pyramidal layer were collected across 2-3 animals per condition. Mounted expanded samples were imaged using an HCX PL APO 40x/1.10 WATER objective. From each dish, 2-4 stacks (0.42 μm Z-step, spanning 250-500 μm across the sample) were collected across 2-3 animals per condition. No deconvolution was used to process any confocal images.

### Stimulated emission depletion (STED) microscopy

Tau-STED super-resolution imaging was performed on a Leica STELLARIS 8 TauSTED 3D FALCON (Leica Microsystems) equipped with a pulsed white light laser for excitation, gated HyD hybrid detectors and 3 lasers for depletion all images were captured with a HC PL APO 100x/1.40 OIL STED W objective. For each fluorophore, excitation wavelength, detection bandwidth and depletion laser were set as recommended by the LASX v4.9.0.30221 software. Multicolor images were acquired in frame sequential mode to avoid cross-excitation and depletion. Only 2D depletion was used. The combination of excitation/depletion lasers were as follows. AlexaFluor 488: 495nm/592nm; AlexaFluor 594+: 593nm/775nm; AlexaFluor 647: 647nm/775nm. For *in vitro* samples, pixel size was set to 30nm with frame average set to 2. For *in vivo* samples, the pixel size was set to 20nm with frame average and line accumulation set to 4. TauSTED detection was applied to *in vivo* samples to enable lifetime-based filtering of fluorescence signals, thus enhancing spatial resolution while maintaining low depletion laser power to prevent excessive bleaching (TauStrength = 400, Denoise = 100). In both *in vitro* and *in vivo* samples, no Z-stacks were taken and no deconvolution was applied. For representative images within this manuscript, FIJI 1.54p was used to subtract the background using a rolling ball radius of 50. For representative images within this manuscript, bilinear interpolation was used to adjust the resolution to >300 dpi.

### Live staining of neuron-astrocyte cocultures with calcein, FM4-64, and PSVue-550

Prior to staining, neuron-astrocyte cocultures were grown and maintained until Day 21. Cultures were incubated with 1 μM Calcein Violet (Invitrogen, Cat #65-0854-39), 4 μM SynaptoRed C2/FM4-64 dye (Biotium, Cat #70021), and/or 5 μM PSVue-550 (Polysciences, Cat #P1005) in fresh neuronal media at 37°C for 20 minutes. After staining, the solution was aspirated and the cells were processed for downstream assays.

When using PSVue dye, 10 mM TES (Sigma-Aldrich, Cat #T1375) was used as a wash buffer. When using SynaptoRed C2/FM4-64 dye, cultures were washed three times for 5 minutes per wash. The second of those washes was supplemented with 500 μM SCAS (Biotium, Cat #70037) to extract cell surface FM4-64 dye. All live stains and associated concentrations are listed in Supplementary Table 2.

### *In vivo* synaptosome preparation

For preparation of synaptosomes from mouse brains, mice were first perfused with 1X PBS on ice before their brains were dissected out. These brains were homogenized in 4 mL of Syn-PER™ Synaptic Protein Extraction Reagent (Thermo Scientific, Catalog #87793) with 30 strokes of a Dounce homogenizer (Genesee Scientific, Cat #09-1853). Homogenate was then centrifuged at 1200xg for 10 minutes at 4°C and the supernatant was transferred to a new tube. This process was repeated three more times. These four pellets were pooled together to make up the cell body fraction which was saved for downstream applications. The remaining supernatant was centrifuged at 15,000xg for 20 minutes at 4°C. The supernatant was aspirated and the remaining pellet formed our synaptosome fraction, which was saved for downstream applications.

### *In vitro* synaptosome preparation

For preparation of synaptosomes from human iPSC-derived neuron-astrocyte cocultures, 300 μL Syn-PER™ Synaptic Protein Extraction Reagent was added to each well. Homogenates were collected by scraping the culture plates and transferred into 1.5 mL microcentrifuge tubes. The homogenate was centrifuged at 1,200xg for 10 minutes at 4°C to remove cell bodies and debris. The supernatant was transferred to a new tube and centrifuged at 15,000xg for 20 minutes at 4°C. The supernatant was aspirated and the remaining pellet formed our synaptosome fraction, which was saved for downstream applications.

### Glycogen Purification

Glycogen (Sigma-Aldrich, Cat #G8751) was dissolved in ultrapure water and an equal volume of Phenol: Chloroform: Isoamyl Alcohol (Invitrogen, Cat #15-593-049) was added. The mixture was centrifuged at 12,000xg for 10 minutes at 4°C to separate the mixture into an aqueous and organic phase. The aqueous phase was transferred to a new tube, and an equal volume of ice-cold chloroform was mixed into the aqueous phase. The mixture was centrifuged again at 12,000xg for 10 minutes at 4°C to separate the mixture into an aqueous and organic phase. An equal volume of 100% ethanol (Sigma-Aldrich, Cat #E7023) was added to the aqueous phase and centrifuged at 12,000xg for 10 minutes at 4°C to generate a purified glycogen pellet. The pellet was allowed to air dry overnight before being dissolved in DNase/RNase-free ultrapure water (Invitrogen, Cat #10977015). Purified glycogen was found to have undetectable levels of RNA contamination as quantified by the Qubit RNA High Sensitivity kit (Invitrogen, Cat #Q32852). Purified glycogen was stored at −80°C till needed.

### RNA extraction

Total RNA was extracted from samples using TRIzol Reagent (Invitrogen, Catalog #15596026) following the manufacturer’s protocols with minor modifications. Samples were homogenized in 1 mL TRIzol with 50 μg of purified glycogen added to facilitate precipitation. 0.2 mL of chloroform was added per 1 mL of TRIzol reagent. Mixtures were centrifuged at 12,000xg for 15 min at 4°C to separate the aqueous phase from the interphase and organic phase. The aqueous phase was collected and RNA was precipitated with 0.5mL of isopropanol (Fisher Scientific, Cat #BP2618-500) per 1 mL of TRIzol reagent. The resulting pellet was washed with 1 mL of fresh 75% ethanol and resuspended in DNase/RNase-Free distilled water. RNA concentration was quantified using the Qubit™ RNA High Sensitivity kit.

### cDNA synthesis

cDNA was synthesized from RNA using the following protocol: RNA samples were digested using ezDNase (Invitrogen, catalog #11766051) for 10 minutes at 37°C. These reactions were converted into cDNA using the Sensifast cDNA Synthesis Kit (Meridian Bioscience, catalog #BIO-65053). cDNA synthesis was performed in a thermocycler following the manufacturer’s recommended PCR cycling protocol.

### Quantitative RT-PCR

Quantitative RT-PCRs were performed using the Sensifast SYBR Lo-Rox kit (Meridian Bioscience, Cat #BIO-94005). Briefly, each reaction was prepared with 1x Sensifast Lo-Rox master mix, 400 nM of the FWD primer, 400 nM of the reverse primer, and 10-100 ng of template cDNA. Quantitative RT-PCR reactions were performed in a Bio-Rad CFX96 Real Time System C1000 Touch Thermocycler running CFX Maestro (version 4.1.24.33.12.19). The thermal cycling conditions consisted of an initial 3 min at 98°C, followed by 39 cycles of 15 sec at 95°C and 20 sec at 60°C followed by 1 sec at 95°C and a 1.92°C s^−1^ ramp from 60°C to 95°C to establish a melting curve. Expression fold changes were calculated using the ΔΔCt method, normalizing to housekeeping gene *GAPDH*. Quantitative RT-PCR primers are listed in Supplementary Table 3.

### Guidecode digital PCR

Digital PCR was performed on a QIAcuity One 5-plex Digital PCR instrument (Qiagen) using 8.5k-partition, 24-well nanoplates (Qiagen, Cat #250011). Digital PCR reactions were prepared with final concentrations of 1X EvaGreen PCR master mix (Qiagen, Cat #250111), 400 nM of the FWD primer, and 400 nM of the REV primer, and 1-100 ng of template cDNA added. Guidecode mRNA level changes were normalized to housekeeping gene *GAPDH*. Digital PCR primer sequences are listed in Supplementary Table 3. The thermal cycling conditions consisted of an initial 2 min at 95°C, followed by 40 cycles of 15 sec at 95°C, 15 sec at 60°C, and 15 sec at 72°C, with a final 5 min cooling step at 40°C. Image capture exposure was set to 250 ms. Data acquisition and analysis were conducted using the QIAcuity 3.1.1 Software Suite. Absolute concentration results from technical replicates were averaged.

### COMPASS-seq guidecode amplicon sequencing

RNA samples were digested using ezDNase for 10 minutes at 37°C to degrade residual DNA. RT-PCR reactions were prepared using a One-Step SSIV RT-PCR Kit (Invitrogen, Cat #1259500) with a final concentration of 1X DNA Polymerase, 500 nM of FWD primer oMK732, and 500 nM of REV indexing primer with 1 μL of Superscript IV reverse-transcriptase and 1 μg of template RNA added to each reaction.

The thermal cycling conditions consisted of an initial 50 minutes at 55°C followed by 30 cycles of the following: 98°C for 10 seconds, 67°C for 15 seconds, and 72°C for 30 seconds. Reactions were incubated for final elongation at 72°C for 5 minutes. Reactions were then SPRI selected for a target amplicon as previously described^33^. Samples were then sequenced by Next-Generation Sequencing using a NextSeq 2000 (Illumina) and demultiplexed with BCLConvert v4.3.13 on Illumina Basespace. Primer sequences are listed in Supplementary Table 3.

### Synaptosome flow cytometry

Day 21 neuron-astrocyte cocultures were live stained for Calcein Violet, SynaptoRed C2/FM4-64, and PSVue-550 as described above. Synaptosome pellets were prepared as described above. Next, synaptosome pellets were resuspended in ST buffer (320 mM sucrose, 5 mM Tris, pH 7.4) and passed through a 35 μm cell strainer (Falcon, Cat #08-771-23) to remove large aggregates. Synaptosome flow cytometry was performed using either a BD FACSAria Fusion set to a sample flow rate of <2 or a BD LSRFortessa X14 set to the lowest sample flow rate. Both machines collected flow data with BD FACSDiva v.9.0.

As previously described^74^, the event detection threshold was set to detect Calcein Violet such that less than 10 events could be detected per second when collecting data from a Calcein Violet-negative synaptosome sample. As previously described^75^, particles were gated to remove large aggregates based on their forward scatter (FSC-A) vs. side scatter (SSC-A) profile. FM+ synaptosomes were subsequently identified based on their FSC-A vs. FM4-64-PerCP-Cy5.5-A profile with gate boundaries defined by the profile of FM-unstained controls such that 1% of unstained synaptosomes fell within the FM+ gate. The gating strategy is represented in Extended Data Figure 4b and Supplementary Figure 1. Detector voltages were optimized using biological samples to avoid saturation. A minimum of 10,000 Calcein Violet-positive events were recorded per sample. All FACS data was processed in FlowJo v10.10.0 software. Biexponential scaling was applied to all scatter and fluorescence parameters to visualize the full dynamic range. For each sample, the median fluorescent intensity was used as the measure of central tendency.

### Sorted synaptosome fixation for immunofluorescent staining

Synaptosomes were prepared for flow cytometry and detected via flow cytometer on a BD FACSAria Fusion flow cytometer as described above. 100,000 FM+ or FM- particles were sorted with the 4-way high purity mask (0-16-0) into each well of a 96-well plate (Cellvis, Cat #P96-1.5P) precoated with 1 mg/mL PDL. Samples were subsequently stained and imaged using the same methods as those described for *in vitro* immunofluorescent staining.

### Somatic flow cytometry

Day 21 neuron-astrocyte cocultures were live stained with PSVue-550. If neurons were not transfected with a fluorescent nuclear marker to distinguish them from astrocytes as in the case of Figure 2d, neurons were live stained with the neuron-specific NeuroFluor NeuO dye (STEMCELL Tech, Cat #01801) per manufacturer’s instructions. Then, cocultures were dissociated for flow cytometry by incubating cocultures with dissociation solution for 1 hour at 37C. Dissociation solution was prepared by mixing 5 mg of lyophilized papain (Worthington, Cat #LK003176) with 5 mL of HBSS, 500 μL of Turbo DNase buffer, and 50 μL of Turbo DNase (Invitrogen, Cat #AM2239). Somatic flow cytometry was performed using a BD LSRFortessa X14 set to the lowest flow rate. Flow data was collected with BD FACSDiva v.9.0. Detected events were gated to remove debris using the forward scatter (FSC-A) vs. side scatter (SSC-A) plot. Single cells were gated based on the forward scatter area (FSC-A) vs. forward scatter height (FSC-H) plot. Neurons were distinguished from astrocytes based on either the nuclear BFP marker expressed exclusively within the neurons or the NeuO-associated fluorescent signal. The gating strategy is represented in Supplementary Figure 1. Detector voltages were optimized using biological samples to avoid saturation. A minimum of 10,000 events were recorded per sample. All FACS data was processed in FlowJo v10.10.0 software. For each sample, the median fluorescent intensity was used as the measure of central tendency.

### AlphaFold 3 modeling

AlphaFold Server^76^ was used to model putative interactions between the ITPR1 receptor and the cytoplasmic N-terminal domain, C-terminal domain, or first intracellular loop of ANO3. Given the size of the ITPR1 receptor and the symmetry of the receptor given its homotetrameric nature, we fully modelled a single ITPR1 monomer and partially modelled portions of the cytoplasmic regulatory domains and C-terminal calcium pore domains of the remaining three ITPR1 monomers composing the complete homotetrameric receptor. The protein sequences of all ITPR1 monomers (complete and partial) along with the protein sequences of the cytoplasmic domains of ANO3 used for AlphaFold modeling are listed in Supplementary Table 14.

To generate the representative model shown in Figure 3m, a model of the ANO3 homodimer was generated based on homology to the experimentally derived protein structure of calcium-bound ANO6 (PDB ID: 6QP6) using SWISS-Model. The ANO3 homodimer model was aligned to the first intracellular loop of ANO3 within an AlphaFold-generated model of the interaction between the ITPR1 receptor and two copies of the first intracellular loop of ANO3 using the matchmaker function on ChimeraX v1.8^77^ aligning the pair of EEEEETLR residues across the two models.

### RNA sequencing and analysis

RNA samples were sent to Plasmidsaurus for RNA sequencing and initial analysis. Briefly, quality of the fastq files was assessed using FastQC v0.12.1. Reads were then quality filtered using fastp v0.24.0 with poly-X tail trimming, 3’ quality-based tail trimming, a minimum Phred quality score of 15, and a minimum length requirement of 50 bp. Quality-filtered reads were aligned to the reference genome using STAR aligner v2.7.11 with non-canonical splice junction removal and output of unmapped reads, followed by coordinate sorting using samtools v1.22.1. PCR and optical duplicates were removed using UMI-based deduplication with UMIcollapse v1.1.0. Alignment quality metrics, strand specificity, and read distribution across genomic features were assessed using RSeQC v5.0.4 and Qualimap v2.3. Gene-level expression quantification was performed using featureCounts (subread package v2.1.1) with strand-specific counting, multi-mapping read fractional assignment, exons and three prime UTR as the feature identifiers, and grouped by gene_id. Final gene counts were annotated with gene biotype and other metadata extracted from the reference GTF file. Sample-sample correlations for PCA were calculated on normalized counts (TMM, trimmed mean of M-values) using Pearson correlation. Differential expression was done with edgeR v4.0.16 using standard practice including filtering for low-expressed genes with edgeR::filterByExprwith default values. Differentially expressed genes are listed in Supplementary Table 4. Differential expression of the synaptically localized mRNAs within CamK2a+ neurons^78^ <u>is shown in</u> Extended Data Figure 1c.

To generate the correlation matrix shown in Extended Data Figure 1a, we set a minimum mean expression of 5 and a minimum variance 0.05 to focus our analysis on the 1,437 most highly expressed and variable RNA species across all samples. Pearson correlation was calculated via the Scipy v1.15.1 stats.pearsonr method to quantify the similarity between each pair of samples. For gene set enrichment analysis, the list of genes and associated average log_2_(fold changes) in RNA abundance between synaptosomes and cell bodies across all samples was generated. This list was analyzed with StringDB’s functional enrichment module with default settings to identify statistically enriched Gene Ontology Cellular Component terms. The complete set of functional enrichment results are provided in Supplementary Table 6.

### CRISPR screen analysis

CRISPR screen analysis was performed as previously described with minor modifications^33^. Briefly, differential sgRNA abundance analysis was performed between synaptosome and cell body samples from each individual mouse. Across all samples, sgRNA abundances were total count normalized.

We excluded any sgRNAs that were represented with fewer than 100 reads within the cell body sample. Guide-level phenotype scores were first calculated for each individual mouse before being pooled together by taking the arithmetic mean across mice. For each mouse, guide-level phenotype scores were defined as the log_2_ transformation of the ratio of the sgRNA abundance from the synaptosome sample divided by the sgRNA abundance from the associated cell body sample from that same mouse.

To calculate guide-level significance scores, a weighted linear regression was used to fit the log variance to the log mean of the cell body sample from a single mouse. The fit variance and mean were then used to parameterize negative binomial distributions for each sgRNA, and a survival function or cumulative distribution function is used to calculate a P value for sgRNA underabundance and overabundance for each mouse. Underabundance and overabundance significance scores for each sgRNA are calculated by taking the geometric mean of the associated underabundance and overabundance p-values calculated from each mouse.

Gene-level phenotype scores were calculated by taking the arithmetic mean of the guide-level phenotype scores for all sgRNAs targeting that gene. As previously described^33^, we created random groupings of non-targeting control guides, which we refer to as amalgam genes. We performed identical calculation as above for each of the newly created amalgam genes.

Gene-level underabundance and overabundance significance scores were calculated for both targeted and amalgam genes by calculating the -log_10_ transformation of the geometric mean of the underabundance and overabundance significance scores for all sgRNAs targeting that gene or composing that specific amalgam gene. Gene level underabundance and overabundance significance scores were normalized to the maximal amalgam gene-associated underabundance and overabundance significance scores. For every gene and amalgam gene, if the gene-level phenotype score was greater than 0, then the gene-level significance score was defined as the gene-level overabundance significance score. Otherwise, the gene-level significance score was defined as the gene-level underabundance significance score. Gene scores were calculated for each gene and amalgam gene by multiplying the gene-level phenotype score by the final gene-level significance score. For each gene, an empirical p-value was calculated by comparing the gene score to the distribution of amalgam gene scores. A Benjamini-Hochberg correction was applied to correct for multiple comparisons and identify hits with a false discovery rate threshold of 10%. Processed CRISPR screen data is presented in Supplementary Table 8.

For visualization on the volcano plot, empirical p-values of 0 were replaced with the p-value of the maximal underabundant or overabundant amalgam gene multiplied by the ratio of the maximal underabundant or overabundant amalgam gene score divided by the gene score of the target gene. For the heatmap visualization shown in Figure 1f, gene level phenotype scores were calculated for each individual mouse and then normalized by the standard deviation of the phenotype scores within that mouse. Processed per-mouse normalized knockdown phenotype data is presented in Supplementary Table 9.

### Synapse count imaging analysis

Cell Profiler 4.2.6 was used to identify Homer and Bassoon object masks from confocal images using the Otsu thresholding method. Colocalization events were identified from the overlap between the two masks. The full pipeline is available at https://kampmannlab.ucsf.edu/shroff-et-al-2026-code.

### Expansion factor approximation

The dense-projection – post-synaptic density (DP-PSD) distance was measured as previously reported^37^. The DP-PSD linear expansion factor was calculated by dividing the mean measured DP-PSD distance by the previously reported DP-PSD distance of 139 nm^79^ as measured by electron microscopy. Raw measurements are provided in Supplementary Table 11.

The nuclear cross-sectional area expansion factor was measured by comparing nuclear cross-sectional area in unexpanded and expanded tissue sections. Expanded nuclei were segmented from using a custom Python script. Bright chromocenters were suppressed by intensity clipping, median filtering, log compression, and Gaussian smoothing before thresholding. Touching nuclei were separated by distance-transform watershed in XY, followed by overlap-based linking across Z slices. Individual nuclei were measured at the middle plane of segmentation mask. Objects below 10,000 pixels or with circularity <0.65 were excluded.

Unexpanded images were first downsampled without interpolation (XY downsampling factor = 0.2; Z unchanged) and median filtered (radius = 5 pixels). Nuclei were segmented using the TrackMate v7.14.0^80^ -StarDist 2D workflow in FIJI. 3D image stacks were treated as pseudo-time series and segmented in 2D on a slice-by-slice basis. Resulting 2D masks were filtered based on morphological and intensity-based features: quality > 0.72, signal-to-noise ratio > 2.32, shape index < 3.85, estimated radius between 3.36 and 4.82 µm, circularity > 0.75, and area > 43.02 µm². Filtered masks were subsequently reconstructed into 3D nuclei using LAP (Linear Assignment Problem) tracking across adjacent Z slices, with a maximum frame-to-frame linking distance of 2.0 µm, maximum gap-closing distance of 1.0 µm, and maximum frame gap of 1 slice. Final 3D masks smaller than 350,000 voxels were excluded.

The mean nuclear cross-sectional area ratio between expanded and unexpanded nuclei was used to calculate the nuclear cross-sectional linear expansion factor as the square root of the area expansion factor. Raw measurements are provided in Supplementary Table 11.

### Nearest neighbor analysis of expansion microscopy immunostaining

Expansion microscopy imaging datasets were analyzed in Fiji using the 3D Object Counter on GPU ( CLIJx v0.32.1.1)^81^ tool with a minimum object size of 5 pixels to identify the 3D coordinates of each punctum across all channels. A custom script was used to analyze these puncta lists to determine the Euclidean centroid-centroid nearest neighbor distance between the closest puncta across distinct fluorescent channels. For the scrambled value datasets used to approximate the distribution of nearest neighbor distances in the case of random puncta, the x and y coordinates are switched for only one of the fluorescent channels.

To approximate the degree of clustering across two fluorescent channels, the probability distribution of nearest-neighbor distances for each punctum in the scrambled data is subtracted from the probability distribution of nearest-neighbor distances for each punctum in the experimental data. A gaussian curve is fit to the first residual peak to approximate the degree of clustering. Amplitudes, mean distances, and standard deviations of these fit curves are reported in Supplementary Tables 12-13. The area under the gaussian curve is calculated by multiplying the amplitude by the standard deviation and 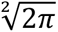 and is reported as the fraction of puncta clustered with the second fluorescent channel.

### Statistical analysis and reproducibility

All statistical analysis unless stated otherwise was performed in GraphPad Prism 11.0.0. Power analyses were performed on preliminary data to determine suitable sample sizes to ensure that experimental power was 80% or greater. All measurements were taken from distinct samples.

Numbers and types of replicates are stated in figure legends. Data were assumed to be normally distributed except for in the case of sgRNA overabundance and underabundance p-values calculated with the assumption of a negative binomial distribution as described in the CRISPR screen analysis section above. For the CRISPR screen, a Benjamini-Hochberg correction was applied to adjust for multiple comparisons. For all other instances where a multiple comparisons correction is appropriate, a Holm-Šídák correction is applied. Data collection and analysis were not performed in a blinded manner to the conditions of the experiments. No animals or data points were excluded from the relevant analyses. Major findings were validated using independent samples and orthogonal approaches.

## Acknowledgements

We thank Carter Brand and Michelle Chang for contributions to preliminary experiments. We thank Julianne Jin, Lin Yadanar, Layla Gomes, and Jaime Leong for general lab support. We thank all members of the Kampmann Lab for advice and technical support. We thank Anna Molofsky, Robert Edwards, Jesse Hanson and Ming-Chi Tsai for discussions. We thank Caroline Mrejen and Alyssa Bonillas (UCSF Weill Advanced Light Microscopy Core) for fluorescent microscopy support. We thank Sarah Elmes, Danielle Peterson, and Christopher Peterson for FACS support.

## Author contributions

K.S. and M.K. conceptualized and led the overall project and wrote the manuscript, with input from all co-authors. K.S., T.J.S., and I.V.L.R designed and developed the *in vivo* CRISPR screening platform for studying compartmental phenotypes. S.I.L, T.J.S., K.S., and M.C.O conducted and analyzed the *in vivo* CRISPRi screen. K.S. designed and developed the neuron-astrocyte coculture model and the FM-dye based synaptosome flow cytometry assay. K.S., T.J.S., and J.F. conducted and analyzed all RNA sequencing, flow cytometry, multielectrode array, and *in vitro* confocal imaging experiments. M.C.O. and K.S.Y. performed all *in vivo* confocal imaging experiments. M.C.O., K.S., and K.S.Y. conducted and analyzed all STED microscopy experiments. M.C.O, K.S., N.R.L., M.E.P., T.J.S., and J.F. conducted and analyzed all expansion microscopy experiments. T.J.S., K.S., and G.A.M. designed and developed the V5-tagged ANO3 iPSC line. S.F., C.C. and L.Y.J. designed and developed the FLAG-GFP11-tagged ANO3 mouse line. K.S. performed all AlphaFold structural interaction modeling.

## Competing interest declaration

MK is a co-scientific founder of Montara Therapeutics and serves on the Scientific Advisory Boards of Montara Therapeutics, Engine Biosciences and Alector, and is an advisor to Modulo Bio and Theseus Therapies. MK is an inventor on US Patent 11,254,933 related to CRISPRi and CRISPRa screening, and on a US Patent application on *in vivo* screening methods.

Supplementary Information is available for this paper.

## Data availability statement

CRISPR screen data will be made available on CRISPRbrain. Synaptosome fractionation RNA sequencing data will be shared on GEO. Expansion microscopy raw data are available upon reasonable request. All other data are available on Dryad.

## Code availability statement

All code and analysis pipelines are available at https://kampmannlab.ucsf.edu/shroff-et-al-2026-code.

## Extended Data Figures/Tables

**Extended Data Figure 1.**
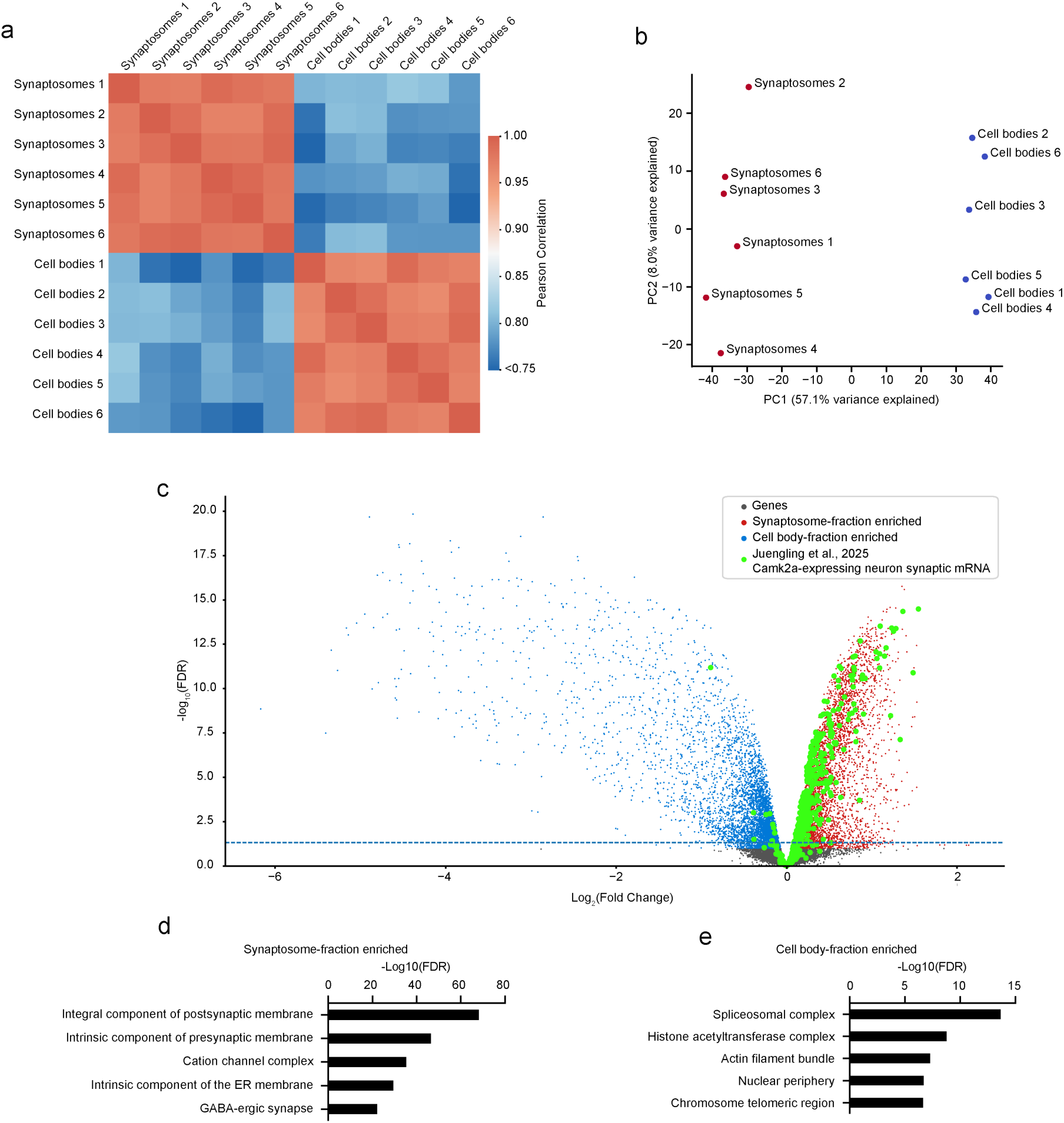
Synaptosome fractionation consistently generates samples enriched for synaptic mRNAs. Six adult P30 mouse brains were collected and fractionated into synaptosome and cell body fractions prior to RNA extraction and sequencing. (a) Pearson correlation analysis revealed that transcriptomes were highly concordant within sample type (synaptosomes, cell bodies), but differed between sample types. (b) PCA analysis shows strong clustering of synaptosome fraction transcriptomes which separated clearly from cell body fraction transcriptomes. (c) Differential gene expression analysis reveals that the synaptosome fractions are robustly enriched for the majority of the 512 synaptically localized mRNAs found in CamK2a+ neurons described in Juengling et al., 2025^72^. The 512 synaptically localized mRNAs found in CamK2a+ neurons are listed in Supplementary Table 5. (d-e) Gene set enrichment analysis of cell component terms reveals that (d) transcripts from both annotated presynaptic and postsynaptic genes are enriched within the synaptosome fractions, while (e) transcripts from genes annotated with nuclear localizations are enriched within the cell body fractions.

**Extended Data Figure 2.**
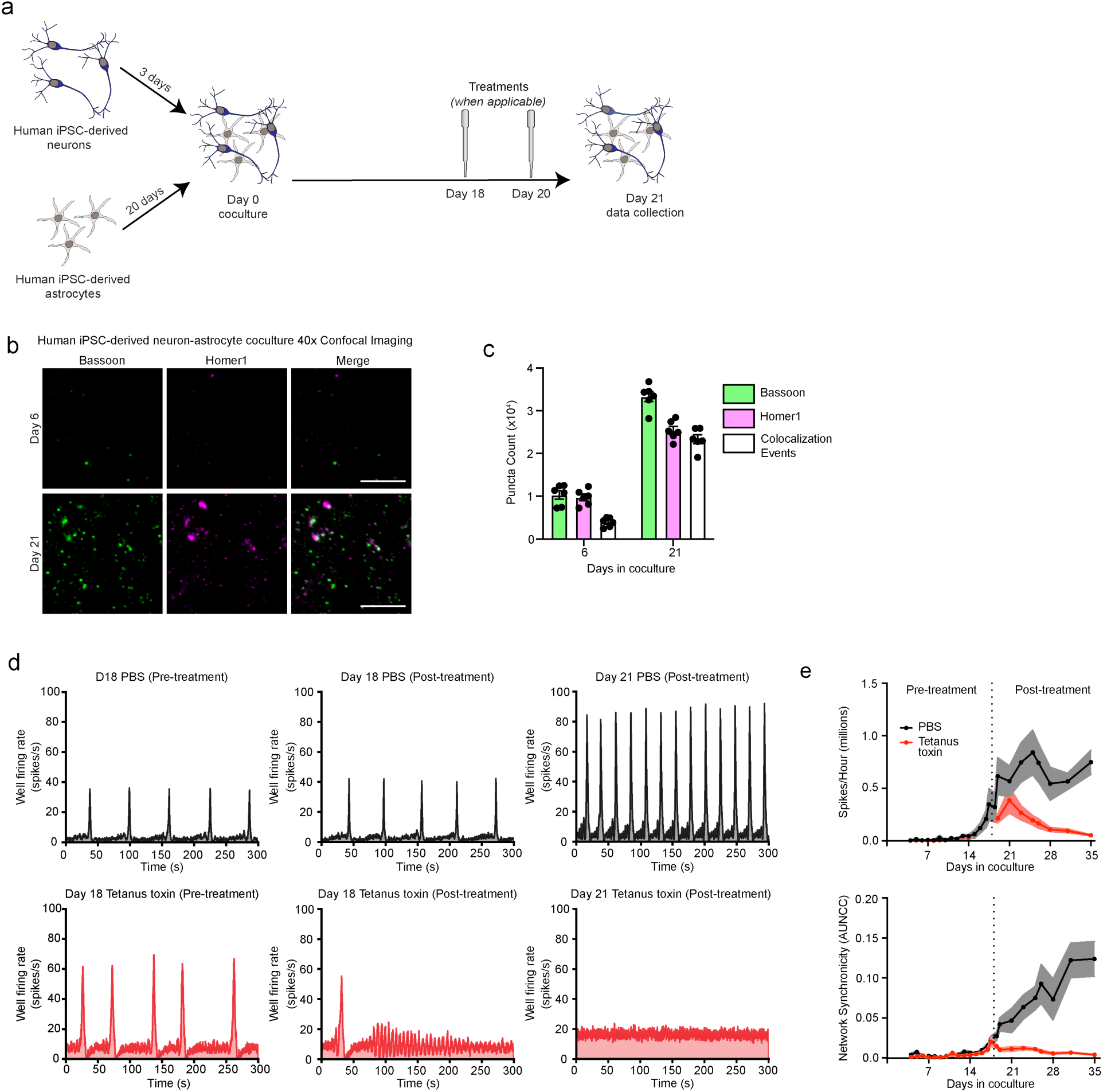
An *in vitro* human iPSC-derived neuron astrocyte coculture model featuring active synapses. (a) Schematic of human iPSC-derived neuron-astrocyte coculture system. (b) Immunofluorescent staining for presynaptic (Bassoon) and postsynaptic (Homer1) markers on Day 6 and Day 21 of coculture maturation (Scale bar = 10 μm). (c) Quantification of structural synapse development across coculture maturation showed a ∼3-fold increase in Bassoon and Homer1 puncta between Day 6 and Day 21 and a ∼6-fold increase in colocalization events between Homer1 and Bassoon puncta (n=6 wells). (d-e) Human iPSC-derived neuron-astrocyte cocultures produce functional synapses contributing to network activity. (d) Representative multi-electrode array traces (left) before treatment with 10 μM tetanus toxin or PBS control, (middle) immediately after treatment with 10 μM tetanus toxin or PBS control, or (right) 72 hours after treatment with 10 μM tetanus toxin or PBS control. (e) Quantification of (top) network spiking activity or (bottom) network synchronicity across coculture maturation when treated on day 18 with either 10 μM tetanus toxin or PBS control (n=6 wells). The solid line represents the mean value while the shaded regions represent the standard error of the mean.

**Extended Data Figure 3.**
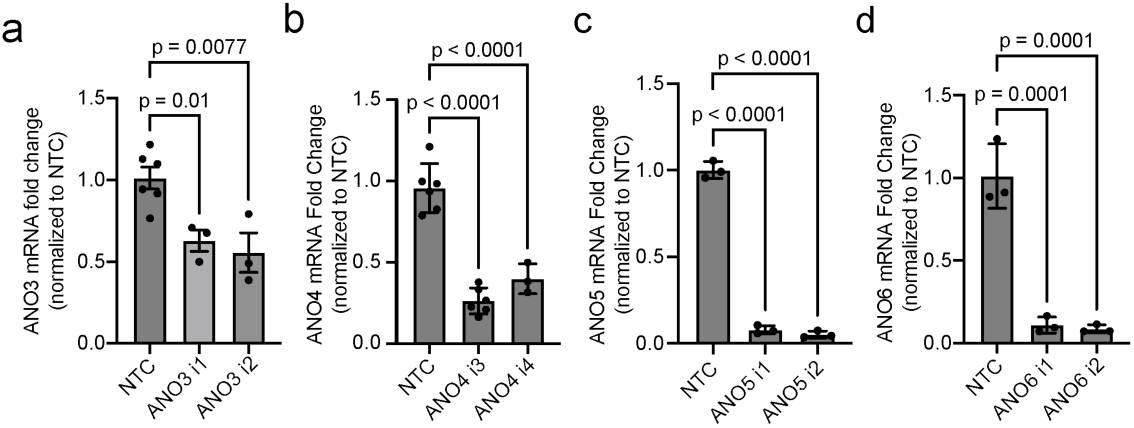
Knockdown of neuronally expressed phospholipid scrambling anoctamins in human iPSCs or human iPSC-derived models. (a-d) Quantification of CRISPRi-mediated knockdown efficiency of sgRNAs targeting neuronally-expressed phospholipid scrambling anoctamins (a) ANO3, (b) ANO4, (c) ANO5, and (d) ANO6 by quantitative RT-PCR (mean ± s.e.m., n = 3-6 technical replicates, p-values calculated by one-way unpaired ANOVA followed by two-sided unpaired t-tests with a Holm-Šídák correction for multiple comparisons applied). Knockdown of ANO3, ANO5, and ANO6 was assessed in human iPSCs. Knockdown of ANO4 was assessed in human iPSC-derived neurons.

**Extended Data Figure 4.**
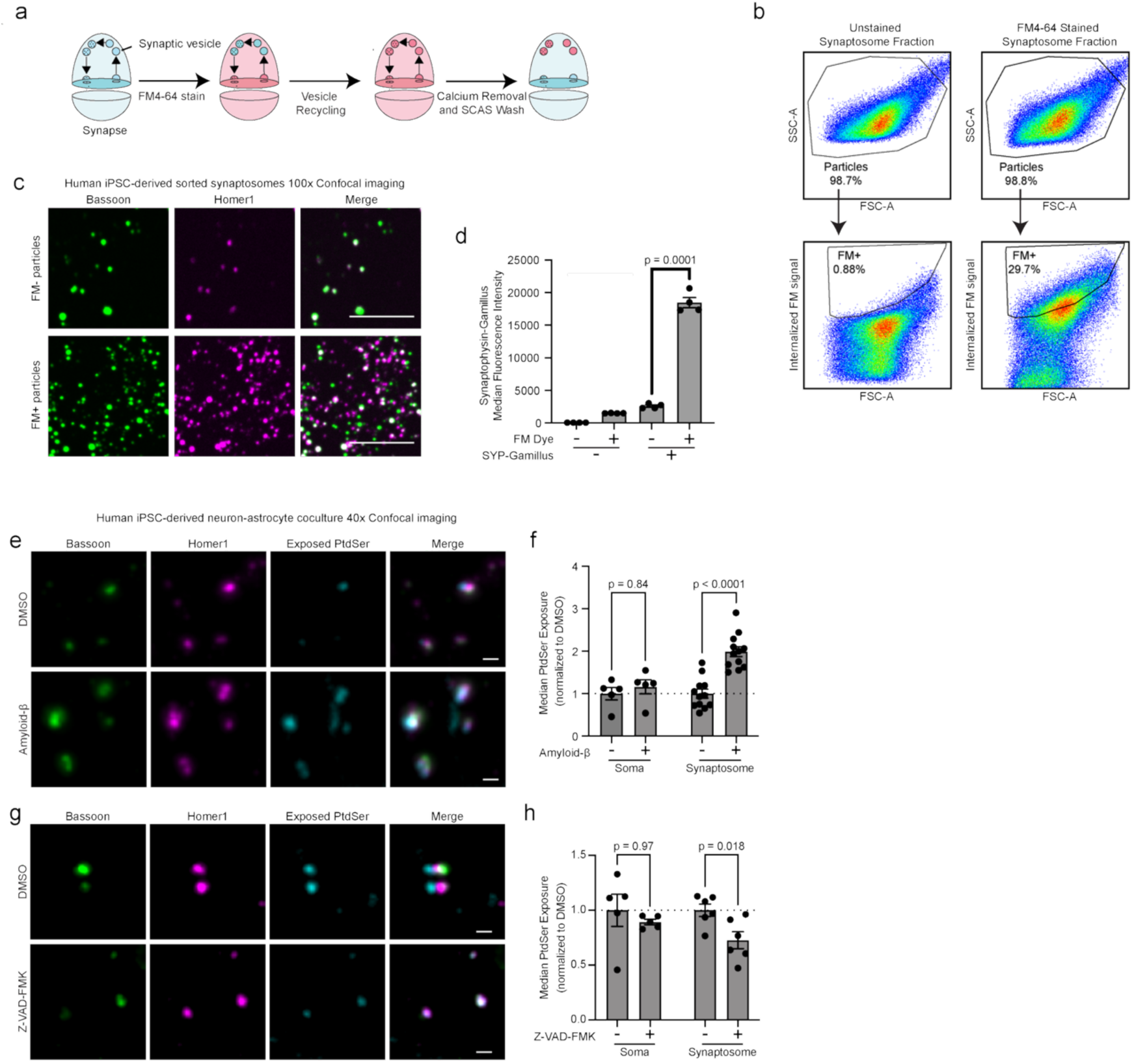
A synaptosome flow cytometry assay to quantify synaptic PtdSer exposure. (a) Schematic of fluorescently labeling of active synapses with FM. (b) Representative synaptosome flow cytometry scatter plots to showcase gating strategy for synaptic particle detection via FM4-64 staining. Events are first detected via fluorescent calcein signaling as recommended in prior literature^79^ on synaptosome flow cytometry. These events are gated (“Particles” gate) on FSC/SSC to remove large aggregates as previously recommended by prior literature^74^. Finally, FM+ particles are identified within experimental samples by drawing the “FM+” gate such that <1% of particles from the FM unstained control fall within the FM+ gate. (c) Immunofluorescent staining of sorted FM+ and FM- particles for presynaptic (Bassoon) and postsynaptic (Homer1) markers (Scale bar = 10 μm). (d) FM+ particles generated from human iPSC-derived neuron-astrocyte cocultures are preferentially enriched ∼7-fold for green fluorescent protein (Gamillus)-labelled synaptophysin relative to FM- particles (mean ± s.e.m., n=4 wells, p-value calculated by two-sided paired t-test). (e,g) Immunofluorescent staining for presynaptic (Bassoon) and postsynaptic (Homer1) markers alongside exposed phosphatidylserine (PS) via pre-fixation Annexin-V staining in human iPSC-derived neurons cocultured with human iPSC-derived astrocytes for 21 days following DMSO control or 72 hours of (e) 10 μg/mL amyloid-β oligomer treatment or (g) 20 μM pan-caspase inhibitor Z-VAD-FMK treatment (Scale bar = 1 μm). (f,h) Flow cytometry reveals that (f) 10 μg/mL amyloid-β oligomer treatment selectively increases synaptic PtdSer exposure, while (h) 20 μM pan-caspase inhibitor Z-VAD-FMK treatment selectively decreases synaptic PtdSer exposure (mean ± s.e.m., n=5-12 wells, p-values calculated by one-way unpaired ANOVA followed by two-sided unpaired t-tests comparing against DMSO control with a Holm-Šídák correction for multiple comparisons applied).

**Extended Data Figure 5.**
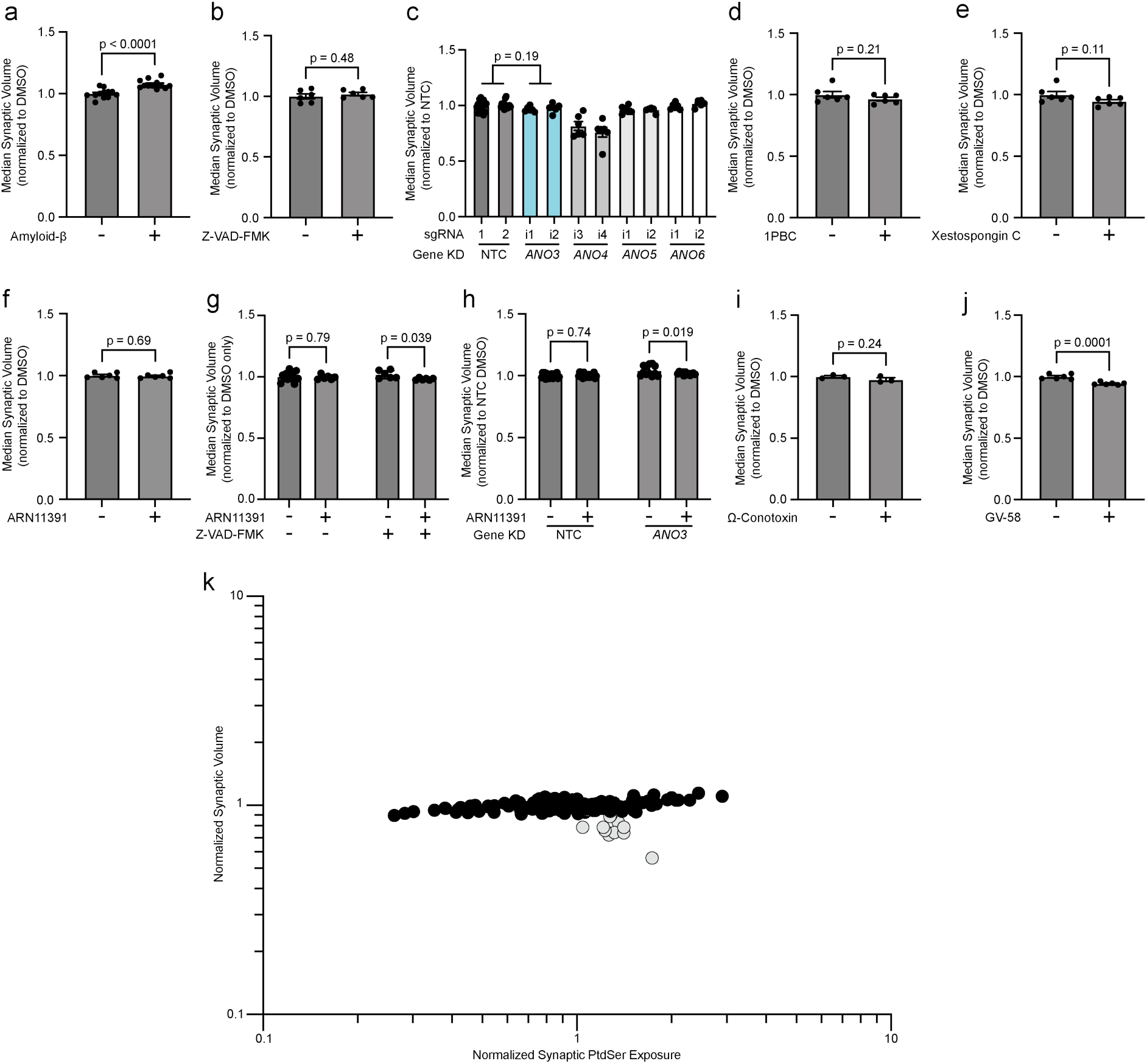
Genetic and pharmacological perturbations that affect synaptic PtdSer exposure negligibly affect synaptosome volume. (a) 72-hour treatment of 10 μg/mL amyloid-β oligomer treatment increases synaptosome volume by ∼7%. (b,d-f,i) 72-hour treatment of (b) 20 μM pan-caspase inhibitor Z-VAD-FMK, (d) 50 μM pan-anoctamin inhibitor 1PBC, (e) 10 μM IP3 receptor inhibitor Xestospongin C, (f) 20 μM ITPR1 activator ARN11391, and (i) 2 μM voltage-gated presynaptic N-type calcium channel inhibitor 0-conotoxin do not affect synaptic volume. (c) CRISPRi-mediated knockdown of ANO3 does not affect synaptosome volume. (g-h) 72-hour treatment of 20 μM ARN11391when combined with either (g) constitutive CRISPRi-mediated ANO3 knockdown or (h) 72 hours of 20 μM pan-caspase inhibitor Z-VAD-FMK treatment reduces synaptosome volume by ∼2%. (j) 72-hour treatment of 50 μM presynaptic voltage-gated calcium channel activator GV-58 decreases synaptosome volume by ∼6% relative to the ∼30% increase in synaptic PtdSer exposure from the same samples (mean ± s.e.m., n= 3-12 wells, p-values calculated by one-way unpaired ANOVA followed by two-sided unpaired t-tests comparing against DMSO or NTC control with a Holm-Šídák correction for multiple comparisons applied). (k) Changes in synaptosome volume negligibly correlate with synaptic PtdSer exposure, apart from a negative correlation within samples where ANO4 expression is knocked down (gray dots).

**Extended Data Figure 6.**
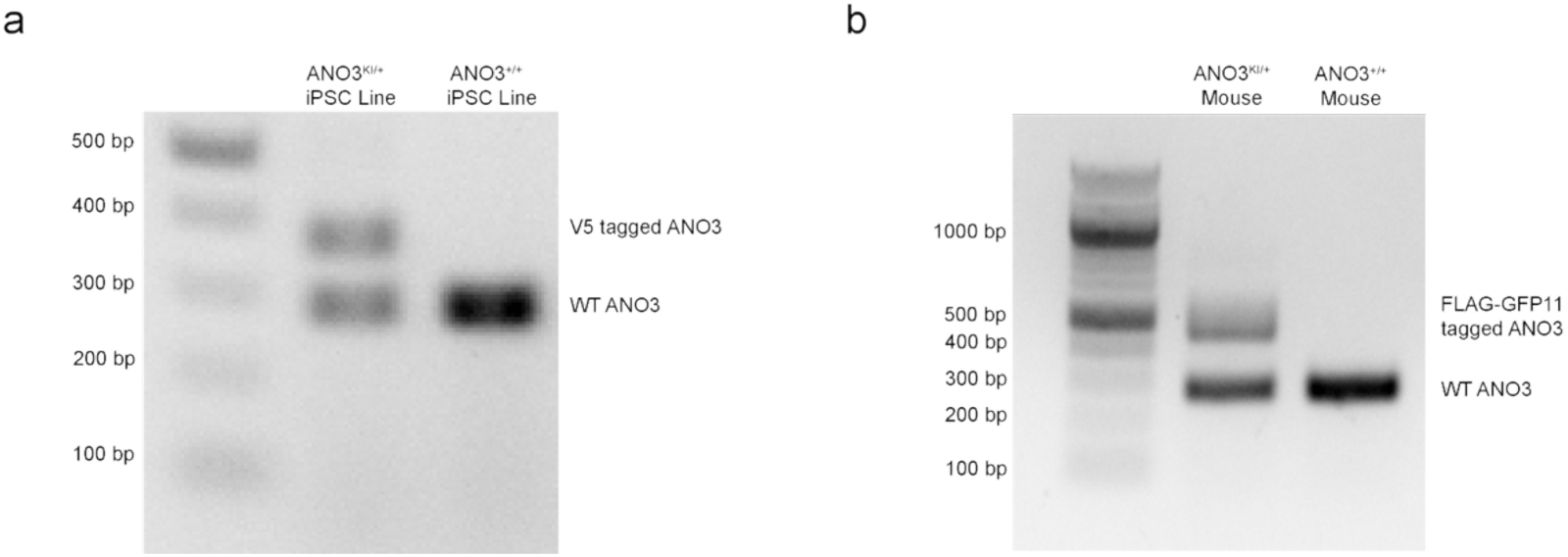
Validation of CRISPR-mediated endogenous tagging of ANO3. (a) Genotyping PCR to assess for successful CRISPR-mediated knock-in of the V5 tag at the N-terminus of ANO3 within human iPSCs. The V5-tagged allele was verified by amplicon sequencing. (b) Genotyping PCR to assess for successful CRISPR-mediated knock-in of the FLAG-GFP11 tag at the C-terminus of ANO3 within mice. The FLAG-GFP11-tagged allele was verified by amplicon sequencing.

**Extended Data Figure 7.**
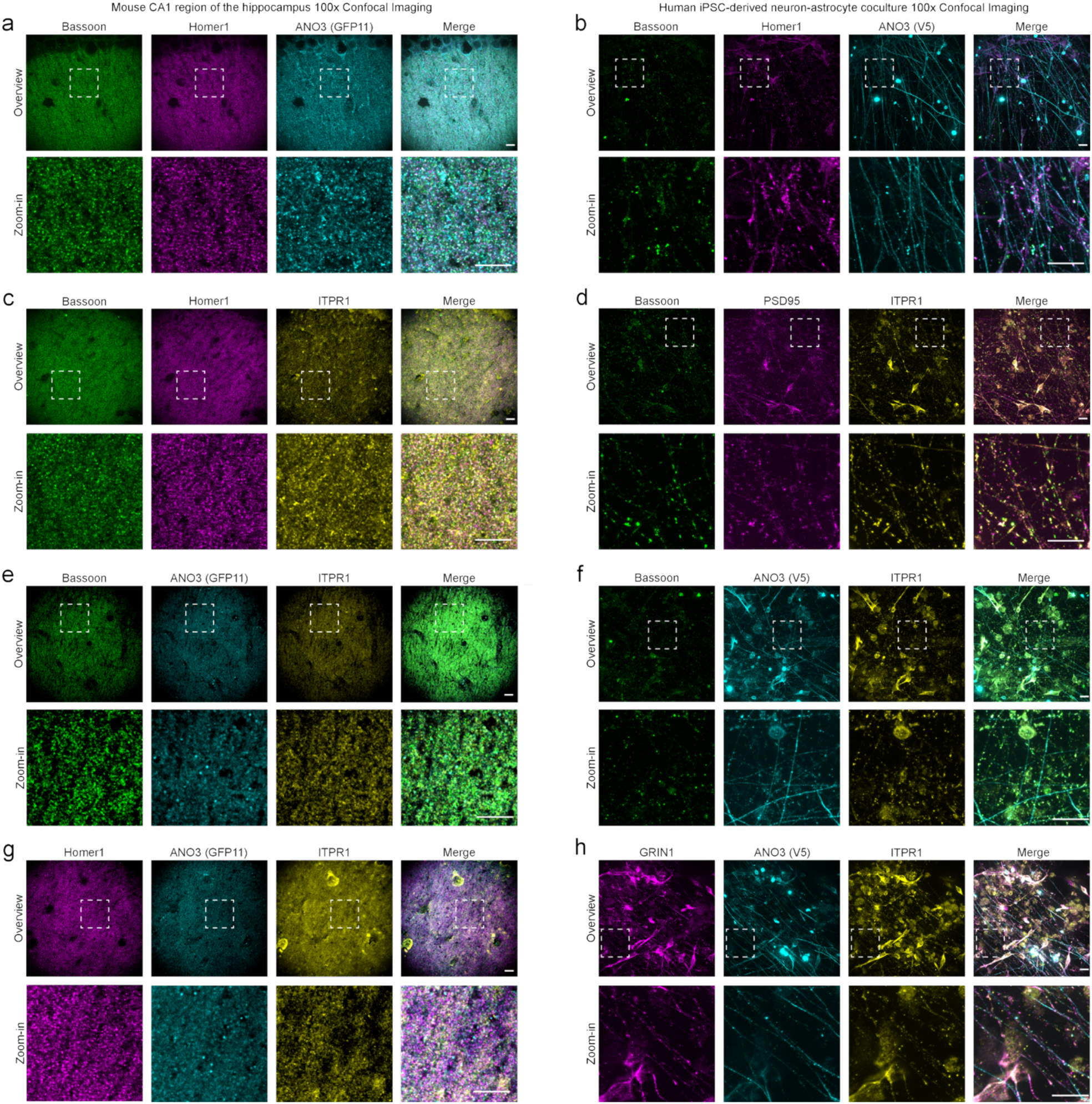
Large fields of view localizing ANO3, ITPR1, and ANO3-ITPR1 clusters to synapses. Immunofluorescent staining and imaging with high-resolution confocal microscopy reveals synaptic localization of (a-b) ANO3, (c-d) ITPR1, and (e-h) ANO3-ITPR1 clusters in the (a,c,e,g) adult mouse CA1 region of the hippocampus and (b,d,f,h) human iPSC-derived neuron-astrocyte cocultures. White dashed box denotes zoomed in region shown in the lower section of each subpanel (Scale bar = 10 μm).

**Extended Data Figure 8.**
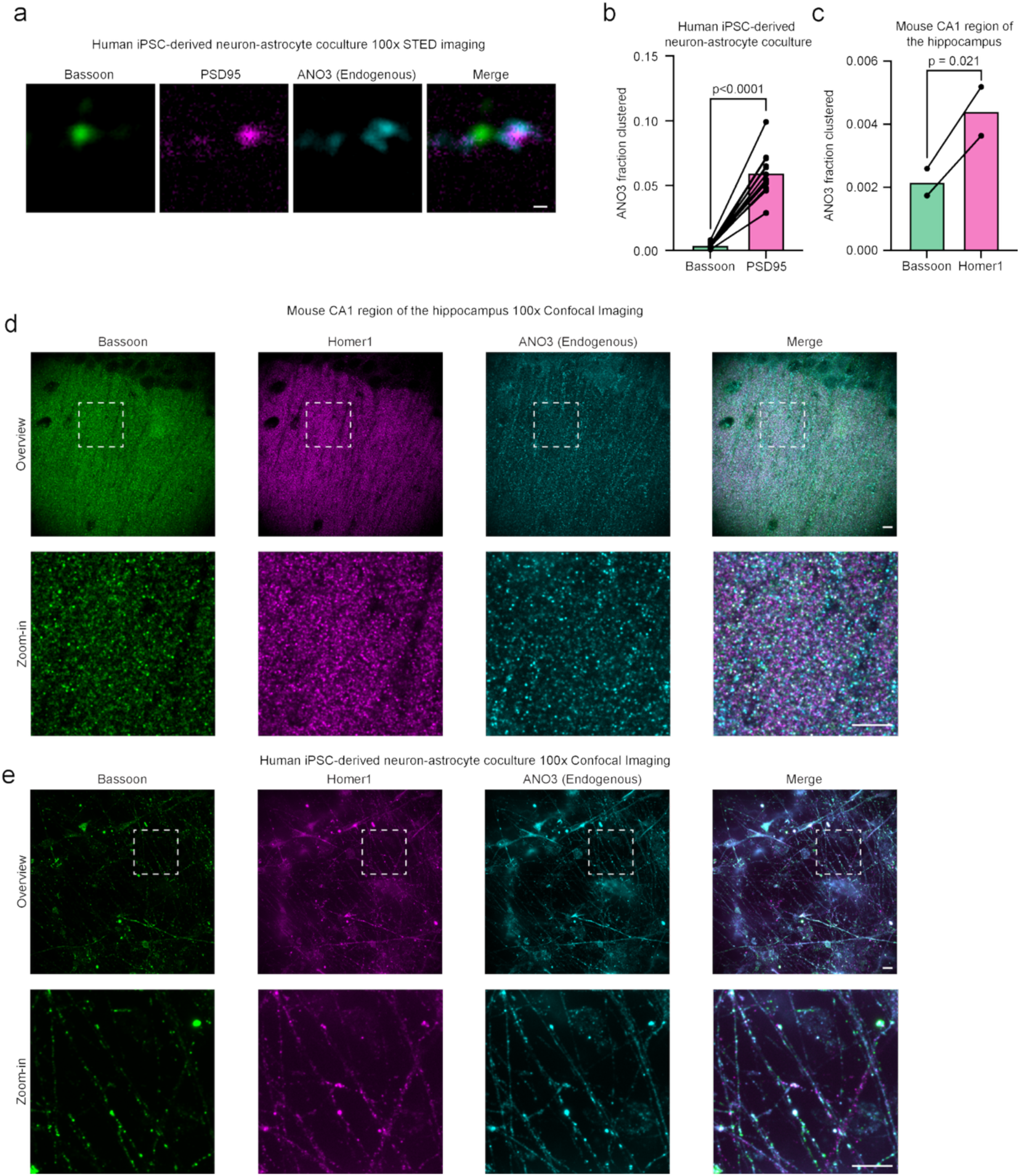
Endogenous ANO3 antibody confirms postsynaptic localization of ANO3. (a) Immunofluorescent staining and imaging with super-resolution STED microscopy reveals that endogenous ANO3 colocalizes to a greater extent with the postsynaptic marker PSD95 than with the presynaptic marker Bassoon in human iPSC-derived neuron-astrocyte cocultures. Representative images are shown here (Scale bar = 200 nm). (b) Quantification of the fraction of ANO3 puncta that colocalize with either Bassoon or PSD95 in iPSC-derived neuron-astrocyte cocultures (mean ± s.e.m., n=12 fields of view across 3 wells collected across 3 sample processing runs, p-value calculated by two-sided paired ratio t-test). (c) Quantification of the fraction of ANO3 puncta that colocalize with either Bassoon or Homer1 in mouse CA1 hippocampus (mean ± s.e.m., n=2 fields of view in 1 mouse collected in 1 sample processing run, p-value calculated by a two-sided paired ratio t-test). (d-e) Immunofluorescent staining and imaging with confocal microscope reveals synaptic localization of ANO3 (endogenous) in (d) adult mouse CA1 region of the hippocampus and (e) human iPSC-derived neuron-astrocyte cocultures. White dashed box denotes zoomed in region shown in the lower section of each subpanel (Scale bar = 10 μm).

**Extended Data Figure 9.**
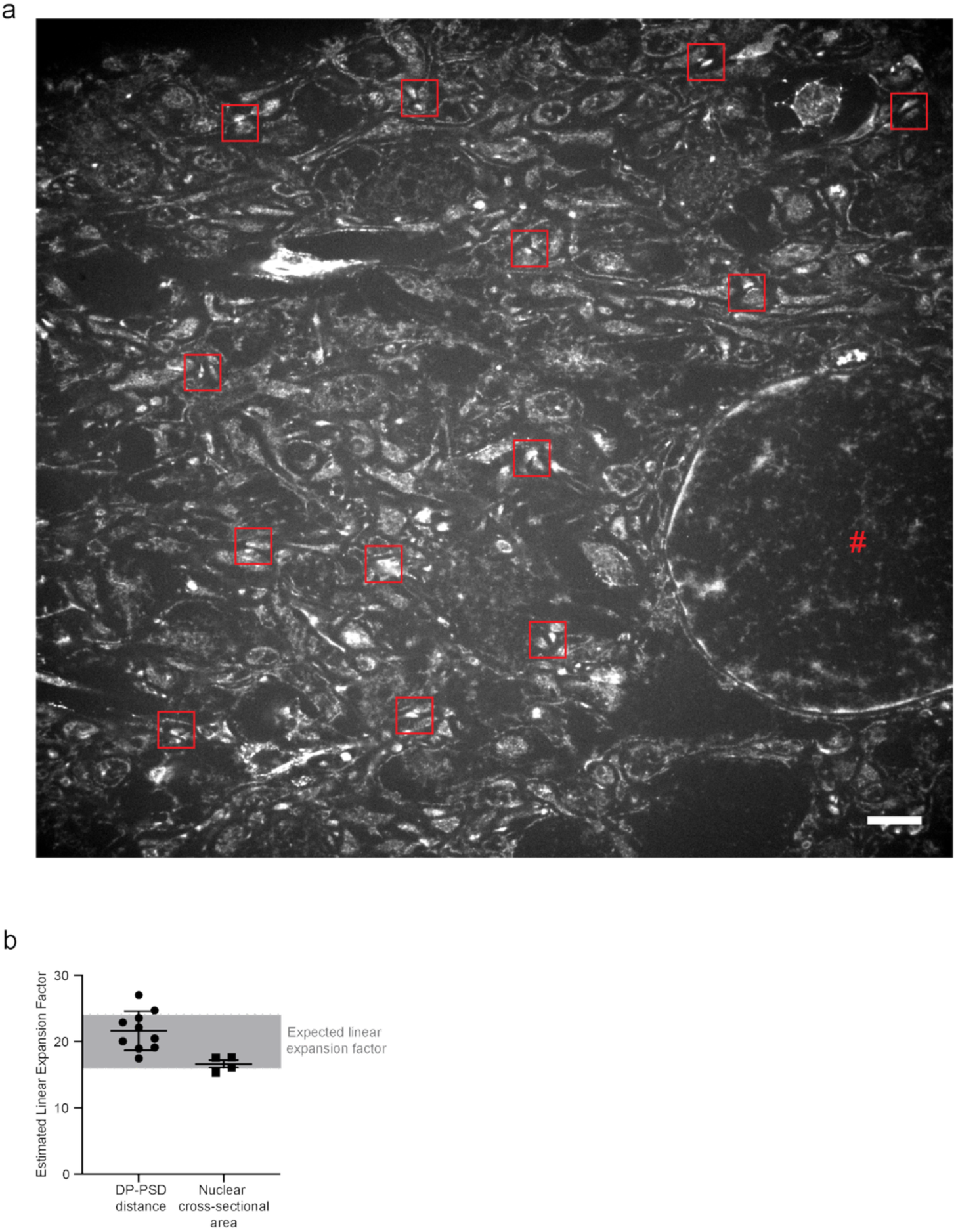
Sample processing for expansion microscopy linearly expands samples by ∼20-fold. (a) Representative total protein stain of expanded mouse CA1 region of the hippocampus tissue reveals expanded nuclei and synapses. Nuclei are denoted by a red hash (#). Synapses have been identified via a red box. (b) (Left) Linear expansion factor approximation using the synaptic dense projection-post synaptic density distance as measured on the total protein stain (mean ± s.d., n= 10 fields of view (250 synapses total) from 4 mice and 4 sample processing runs) normalized to the previously reported unexpanded synaptic dense projection-post synaptic density distance of 139 nm measured via electron microscopy^77^. (Right) Linear expansion factor approximation using the nuclear cross-sectional area as measured via SYTOX Green signal in expanded samples (mean ± s.d., n= 4 fields of view (166 nuclei total) from 4 mice and 4 sample processing runs) normalized to the empirically determined nuclear cross-sectional area of unexpanded nuclei as measured via Hoechst signal (73 ± 9 um^2^; n=99 nuclei across 4 mice).

**Extended Data Figure 10.**
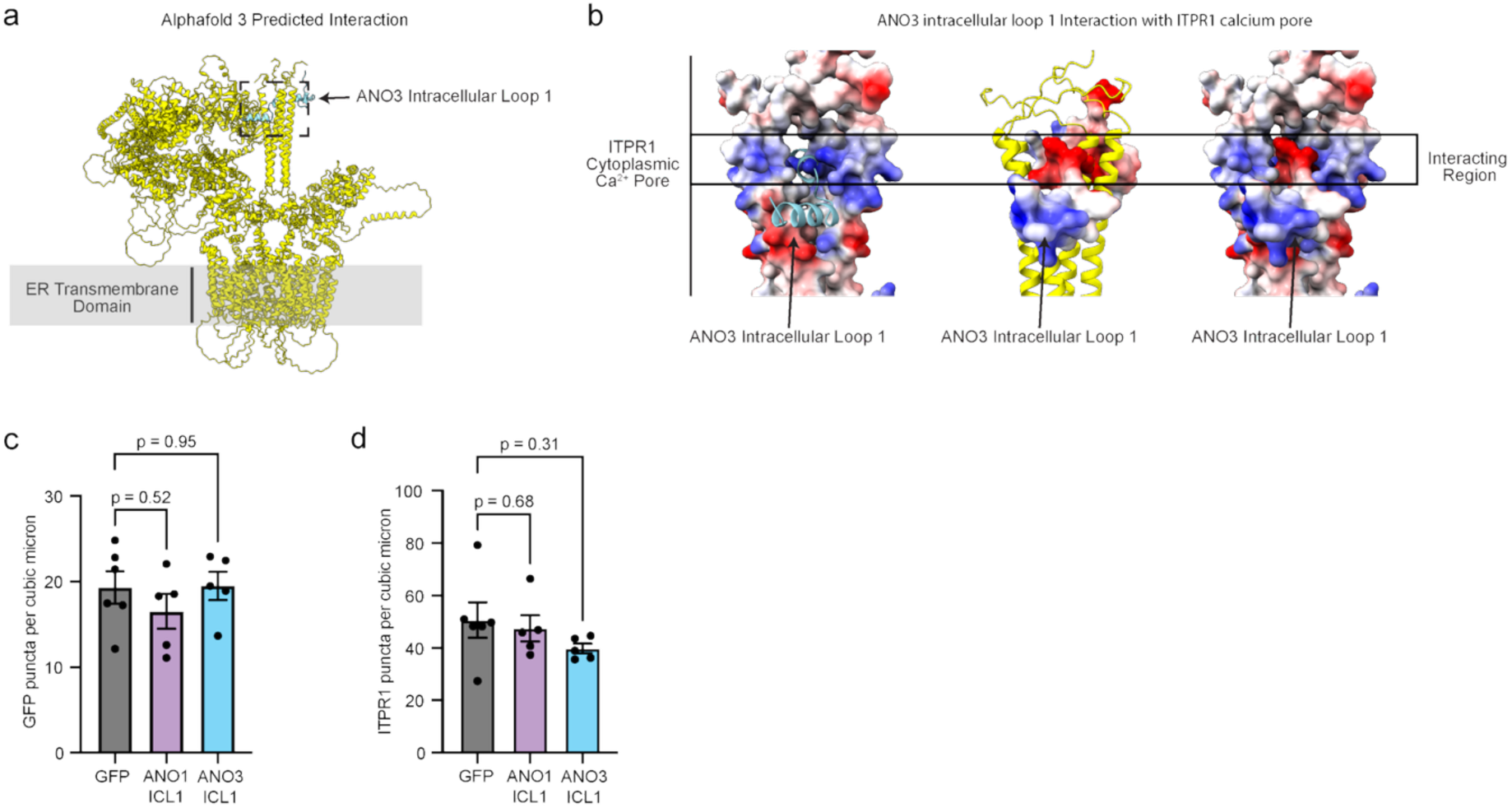
Assessing the predicted interaction between ITPR1 and the first intracellular loop of ANO3. (a) Visualization of the Alphafold 3 predicted interaction between ITPR1 and the first intracellular loop of ANO3. (b) Visualizations of the coulombic charge potentials at the interaction interface where there appears to be a potential electrostatic interaction between the acidic residues on ANO3’s first intracellular loop and the basic residues at the distal end of ITPR1’s C-terminal domain. (c-d) Quantification of the (c) GFP puncta density and the (d) ITPR1 puncta density in the expanded samples analyzed in Figure 3n (mean ± s.e.m., *In vitro*, n = 6 fields of view across 2 wells and 1 sample processing run. *In vivo*, n = 4 fields of view across 1 mouse and 1 sample processing run, p-values calculated by one-way unpaired lognormal ANOVA followed by two-sided unpaired lognormal t-tests with a Holm-Šídák correction for multiple comparisons applied).

**Supplementary Figure 1:**
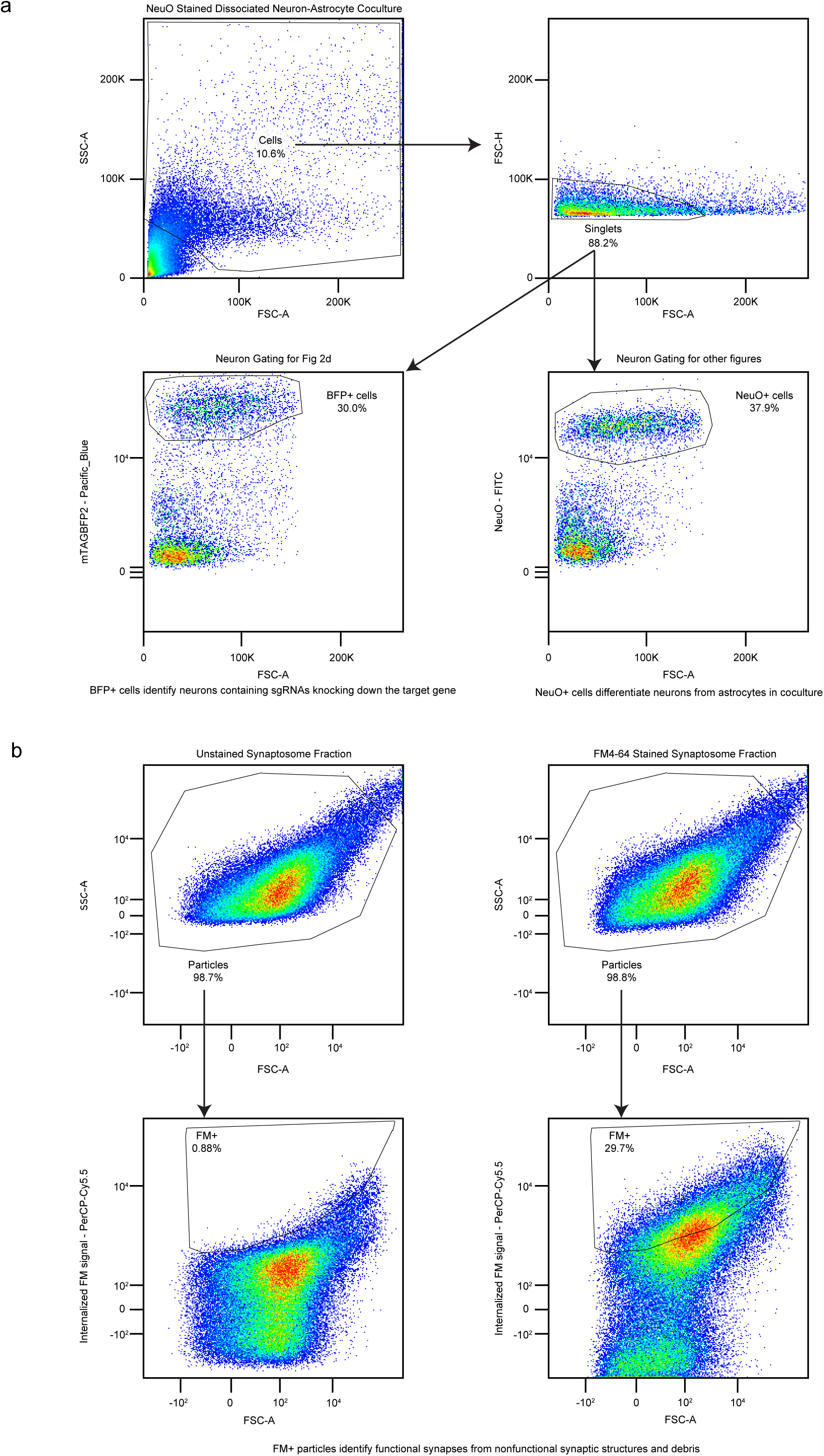
Flow cytometry gating strategies. Representative flow cytometry data highlighting (a) somatic flow cytometry gating strategy and (b) synaptosome flow cytometry gating strategy.

## Main references

1. Suzuki, J., Denning, D. P., Imanishi, E., Horvitz, H. R. & Nagata, S. Xk-Related Protein 8 and CED-8 Promote Phosphatidylserine Exposure in Apoptotic Cells. Science 341, 403–406 (2013).

2. Segawa, K. et al. Caspase-mediated cleavage of phospholipid flippase for apoptotic phosphatidylserine exposure. Science 344, 1164–1168 (2014).

3. Györffy, B. A. et al. Local apoptotic-like mechanisms underlie complement-mediated synaptic pruning. Proc. Natl. Acad. Sci. U. S. A. 115, 6303–6308 (2018).

4. Sokolova, D., Childs, T. & Hong, S. Insight into the role of phosphatidylserine in complement-mediated synapse loss in Alzheimer’s disease. Fac. Rev. 10, 19 (2021).

5. Yu, Z., Gutu, A., Kim, N. & O’Shea, E. K. Activity-dependent synapse elimination requires caspase-3 activation. eLife 13, (2025).

6. Andoh, M. et al. Nonapoptotic caspase-3 guides C1q-dependent synaptic phagocytosis by microglia. Nat. Commun. 16, 918 (2025).

7. Mega Vascular Cognitive Impairment and Dementia (MEGAVCID) consortium. A genome-wide association meta-analysis of all-cause and vascular dementia. Alzheimers Dement. J. Alzheimers Assoc. 20, 5973–5995 (2024).

8. Feenstra, B. et al. Common variants associated with general and MMR vaccine–related febrile seizures. Nat. Genet. 46, 1274–1282 (2014).

9. Charlesworth, G. et al. Mutations in ANO3 Cause Dominant Craniocervical Dystonia: Ion Channel Implicated in Pathogenesis. Am. J. Hum. Genet. 91, 1041–1050 (2012).

10. Katz, L. C. & Shatz, C. J. Synaptic activity and the construction of cortical circuits. Science 274, 1133–1138 (1996).

11. Faust, T., Gunner, G. & Schafer, D. P. Mechanisms governing activity-dependent synaptic pruning in the mammalian CNS. Nat. Rev. Neurosci. 22, 657–673 (2021).

12. Kano, M. & Hashimoto, K. Synapse elimination in the central nervous system. Curr. Opin. Neurobiol. 19, 154–161 (2009).

13. Hua, J. Y. & Smith, S. J. Neural activity and the dynamics of central nervous system development. Nat. Neurosci. 7, 327–332 (2004).

14. Penzes, P., Buonanno, A., Passafarro, M., Sala, C. & Sweet, R. A. Developmental Vulnerability of Synapses and Circuits Associated with Neuropsychiatric Disorders. J. Neurochem. 126, 165–182 (2013).

15. Wiegert, J. S. & Oertner, T. G. Long-term depression triggers the selective elimination of weakly integrated synapses. Proc. Natl. Acad. Sci. U. S. A. 110, E4510–E4519 (2013).

16. Malenka, R. C. & Bear, M. F. LTP and LTD: An Embarrassment of Riches. Neuron 44, 5–21 (2004).

17. Piochon, C., Kano, M. & Hansel, C. LTD-like molecular pathways in developmental synaptic pruning. Nat. Neurosci. 19, 1299–1310 (2016).

18. Schafer, D. P. et al. Microglia sculpt postnatal neural circuits in an activity and complement-dependent manner. Neuron 74, 691–705 (2012).

19. Stevens, B. et al. The classical complement cascade mediates CNS synapse elimination. Cell 131, 1164–1178 (2007).

20. Paolicelli, R. C. et al. Synaptic pruning by microglia is necessary for normal brain development. Science 333, 1456–1458 (2011).

21. Païdassi, H. et al. C1q Binds Phosphatidylserine and Likely Acts as a Multiligand-Bridging Molecule in Apoptotic Cell Recognition. J. Immunol. Baltim. Md 1950 180, 2329–2338 (2008).

22. Scott-Hewitt, N. et al. Local externalization of phosphatidylserine mediates developmental synaptic pruning by microglia. EMBO J. 39, e105380 (2020).

23. Park, D. et al. BAI1 is an engulfment receptor for apoptotic cells upstream of the ELMO/Dock180/Rac module. Nature 450, 430–434 (2007).

24. Chung, W.-S. et al. Astrocytes mediate synapse elimination through MEGF10 and MERTK pathways. Nature 504, 394–400 (2013).

25. Rueda-Carrasco, J. et al. Microglia-synapse engulfment via PtdSer-TREM2 ameliorates neuronal hyperactivity in Alzheimer’s disease models. EMBO J. 42, e113246 (2023).

26. Lemke, G. How macrophages deal with death. Nat. Rev. Immunol. 19, 539–549 (2019).

27. Bevers, E. M. & Williamson, P. L. Getting to the Outer Leaflet: Physiology of Phosphatidylserine Exposure at the Plasma Membrane. Physiol. Rev. 96, 605–645 (2016).

28. Suzuki, J. et al. Calcium-dependent phospholipid scramblase activity of TMEM16 protein family members. J. Biol. Chem. 288, 13305–13316 (2013).

29. Kampmann, M. CRISPRi and CRISPRa screens in mammalian cells for precision biology and medicine. ACS Chem. Biol. 13, 406–416 (2018).

30. Sanjana, N. E. Genome-scale CRISPR pooled screens. Anal. Biochem. 532, 95–99 (2017).

31. Gross, G. G. et al. Recombinant probes for visualizing endogenous synaptic proteins in living neurons. Neuron 78, 971–985 (2013).

32. Cilley, C. D. & Williamson, J. R. Analysis of bacteriophage N protein and peptide binding to boxB RNA using polyacrylamide gel coelectrophoresis (PACE). RNA 3, 57–67 (1997).

33. Ramani, B. et al. CRISPR screening by AAV episome-sequencing (CrAAVe-seq): a scalable cell-type-specific in vivo platform uncovers neuronal essential genes. Nat. Neurosci. 28, 2129–2140 (2025).

34. Neniskyte, U., Neher, J. J. & Brown, G. C. Neuronal Death Induced by Nanomolar Amyloid β Is Mediated by Primary Phagocytosis of Neurons by Microglia. J. Biol. Chem. 286, 39904–39913 (2011).

35. Feng, S. et al. Identification of a drug binding pocket in TMEM16F calcium-activated ion channel and lipid scramblase. Nat. Commun. 14, 4874 (2023).

36. Tillberg, P. W. & Chen, F. Expansion Microscopy: Scalable and Convenient Super-Resolution Microscopy. Annu. Rev. Cell Dev. Biol. 35, 683–701 (2019).

37. M’Saad, O. et al. All-optical visualization of specific molecules in the ultrastructural context of brain tissue. Nat. Biotechnol. 1–15 (2025) doi:10.1038/s41587-025-02905-4.

38. Jin, X. et al. Activation of the Cl- channel ANO1 by localized calcium signals in nociceptive sensory neurons requires coupling with the IP3 receptor. Sci. Signal. 6, ra73 (2013).

39. Inoue, T., Kato, K., Kohda, K. & Mikoshiba, K. Type 1 Inositol 1,4,5-Trisphosphate Receptor Is Required for Induction of Long-Term Depression in Cerebellar Purkinje Neurons. J. Neurosci. 18, 5366–5373 (1998).

40. Khodakhah, K. & Armstrong, C. M. Induction of long-term depression and rebound potentiation by inositol trisphosphate in cerebellar Purkinje neurons. Proc. Natl. Acad. Sci. U. S. A. 94, 14009–14014 (1997).

41. Carrasco, S. & Meyer, T. STIM Proteins and the Endoplasmic Reticulum-Plasma Membrane Junctions. Annu. Rev. Biochem. 80, 973–1000 (2011).

42. Chang, C.-L., Chen, Y.-J. & Liou, J. ER-plasma membrane junctions: Why and how do we study them? Biochim. Biophys. Acta BBA - Mol. Cell Res. 1864, 1494–1506 (2017).

43. Chang, C.-L. et al. Feedback Regulation of Receptor-Induced Ca2+ Signaling Mediated by E-Syt1 and Nir2 at Endoplasmic Reticulum-Plasma Membrane Junctions. Cell Rep. 5, 813–825 (2013).

44. Le, S. C. & Yang, H. An Additional Ca2+ Binding Site Allosterically Controls TMEM16A Activation. Cell Rep. 33, 108570 (2020).

45. Khelashvili, G., Kots, E., Cheng, X., Levine, M. V. & Weinstein, H. The allosteric mechanism leading to an open-groove lipid conductive state of the TMEM16F scramblase. *Commun*. Biol. 5, 990 (2022).

46. Gafni, J. et al. Xestospongins: potent membrane permeable blockers of the inositol 1,4,5-trisphosphate receptor. Neuron 19, 723–733 (1997).

47. Genovese, M. et al. Pharmacological potentiators of the calcium signaling cascade identified by high-throughput screening. PNAS Nexus 2, pgac288 (2023).

48. Nakamura, T., Barbara, J.-G., Nakamura, K. & Ross, W. N. Synergistic Release of Ca2+ from IP3-Sensitive Stores Evoked by Synaptic Activation of mGluRs Paired with Backpropagating Action Potentials. Neuron 24, 727–737 (1999).

49. Kerr, L. M. & Yoshikami, D. A venom peptide with a novel presynaptic blocking action. Nature 308, 282–284 (1984).

50. Tarr, T. B. et al. Evaluation of a novel calcium channel agonist for therapeutic potential in Lambert-Eaton myasthenic syndrome. J. Neurosci. Off. J. Soc. Neurosci. 33, 10559–10567 (2013).

51. Urke, A. et al. Protein-guided RNA barcoding links transcriptomes to synaptic architecture. 2026.02.26.705527 Preprint at 10.64898/2026.02.26.705527 (2026).

52. Yao, Z. et al. A taxonomy of transcriptomic cell types across the isocortex and hippocampal formation. Cell 184, 3222–3241.e26 (2021).

53. Huang, F. et al. TMEM16C facilitates Na(+)-activated K+ currents in rat sensory neurons and regulates pain processing. Nat. Neurosci. 16, 1284–1290 (2013).

54. Um, J. W. et al. Metabotropic Glutamate Receptor 5 is a Co-Receptor for Alzheimer Aβ Oligomer Bound to Cellular Prion Protein. Neuron 79, 887–902 (2013).

55. Stoner, A. et al. Neuronal transcriptome, tau and synapse loss in Alzheimer’s knock-in mice require prion protein. Alzheimers Res. Ther. 15, 201 (2023).

56. Haas, L. T. et al. Metabotropic glutamate receptor 5 couples cellular prion protein to intracellular signalling in Alzheimer’s disease. Brain 139, 526–546 (2016).

57. Ye, W. et al. Phosphatidylinositol-(4, 5)-bisphosphate regulates calcium gating of small-conductance cation channel TMEM16F. Proc. Natl. Acad. Sci. 115, (2018).

58. Yu, K., Jiang, T., Cui, Y., Tajkhorshid, E. & Hartzell, H. C. A network of phosphatidylinositol 4,5-bisphosphate binding sites regulates gating of the Ca2+-activated Cl− channel ANO1 (TMEM16A). Proc. Natl. Acad. Sci. 116, 19952–19962 (2019).

59. Xiao, Q. et al. Voltage- and calcium-dependent gating of TMEM16A/Ano1 chloride channels are physically coupled by the first intracellular loop. Proc. Natl. Acad. Sci. 108, 8891–8896 (2011).

60. Li, T. et al. Phospholipid-flippase chaperone CDC50A is required for synapse maintenance by regulating phosphatidylserine exposure. EMBO J. 40, e107915 (2021).

61. Lehrman, E. K. et al. CD47 Protects Synapses from Excess Microglia-Mediated Pruning during Development. Neuron 100, 120–134.e6 (2018).

## Methods references

62. Mattis, J. et al. Principles for applying optogenetic tools derived from direct comparative analysis of microbial opsins. Nat. Methods 9, 159–172 (2011).

63. Bensussen, S. et al. A Viral Toolbox of Genetically Encoded Fluorescent Synaptic Tags. iScience 23, 101330 (2020).

64. Chen, J. J. et al. Compromised function of the ESCRT pathway promotes endolysosomal escape of tau seeds and propagation of tau aggregation. J. Biol. Chem. 294, 18952–18966 (2019).

65. Peikon, I. D. et al. Using high-throughput barcode sequencing to efficiently map connectomes. Nucleic Acids Res. 45, e115 (2017).

66. Samelson, A. J. et al. Kinetic and structural comparison of a protein’s cotranslational folding and refolding pathways. Sci. Adv. 4, eaas9098 (2018).

67. Tian, R. et al. CRISPR interference-based platform for multimodal genetic screens in human iPSC-derived neurons. Neuron 104, 239–255.e12 (2019).

68. Heo, S.-J. et al. Compact CRISPR genetic screens enabled by improved guide RNA library cloning. Genome Biol. 25, 25 (2024).

69. Chan, K. Y. et al. Engineered AAVs for efficient noninvasive gene delivery to the central and peripheral nervous systems. Nat. Neurosci. 20, 1172–1179 (2017).

70. Miyaoka, Y. et al. Isolation of single-base genome-edited human iPS cells without antibiotic selection. Nat. Methods 11, 291–293 (2014).

71. Leng, K. et al. CRISPRi screens in human iPSC-derived astrocytes elucidate regulators of distinct inflammatory reactive states. Nat. Neurosci. 25, 1528–1542 (2022).

72. Gill, K., Sachdev, A., Conklin, B. & Clelland, C. D. Creating iPSC lines with Ribonucleoprotein (RNP): Nucleofection, Single-cell Sorting, Genotyping, and Line … https://www.protocols.io/view/creating-ipsc-lines-with-ribonucleoprotein-rnp-nuc-4r3l2oeopv1y/v1 (2022).

73. Edelstein, A. D. et al. Advanced methods of microscope control using μManager software. J. Biol. Methods 1, 1 (2014).

74. Hobson, B. D. & Sims, P. A. Critical Analysis of Particle Detection Artifacts in Synaptosome Flow Cytometry. eNeuro 6, ENEURO.0009-19.2019 (2019).

75. Luquet, E., Biesemann, C., Munier, A. & Herzog, E. Purification of Synaptosome Populations Using Fluorescence-Activated Synaptosome Sorting. Methods Mol. Biol. 1538, 121–134 (2017).

76. Abramson, J. et al. Accurate structure prediction of biomolecular interactions with AlphaFold 3. Nature 630, 493–500 (2024).

77. Meng, E. C. et al. UCSF ChimeraX: Tools for structure building and analysis. Protein Sci. Publ. Protein Soc. 32, e4792 (2023).

78. Juengling, M. et al. Brain-wide synaptosome profiling reveals localized mRNAs that diversify synapses. 2025.12.07.692806 Preprint at 10.64898/2025.12.07.692806 (2025).

79. Tavakoli, M. R. et al. Light-microscopy-based connectomic reconstruction of mammalian brain tissue. Nature 642, 398–410 (2025).

80. Ershov, D. et al. TrackMate 7: integrating state-of-the-art segmentation algorithms into tracking pipelines. Nat. Methods 19, 829–832 (2022).

81. Haase, R. et al. CLIJ: GPU-accelerated image processing for everyone. Nat. Methods 17, 5–6 (2020).

